# Genomic diversity of non-typhoidal *Salmonella* found in patients suffering from gastroenteritis in Norfolk, UK

**DOI:** 10.1101/2025.03.06.641842

**Authors:** Steven J Rudder, Bilal Djeghout, Ngozi Elumogo, Nicol Janecko, Gemma C Langridge

**Affiliations:** Quadram Institute Bioscience, Rosalind Franklin Rd, Norwich Research Park, Norwich NR4 7UQ, UK; Centre for Microbial Interaction, Rosalind Franklin Rd, Norwich Research Park, Norwich NR4 7UQ, UK; Eastern Pathology Alliance, Norfolk and Norwich University Hospital, Norwich NR4 7UY, UK

**Keywords:** Non-typhoidal *Salmonella*, genomic diversity, gastroenteritis

## Abstract

*Salmonella* is a significant public health pathogen responsible for a wide spectrum of diseases, ranging from gastroenteritis to invasive non-typhoidal salmonellosis (iNTS) and enteric fever. Although advancements in whole genome sequencing (WGS) have improved surveillance and outbreak investigations, traditional single-colony sequencing methods overlook within-host diversity, potentially underestimating the complexity of infections. This study explores the genome-wide diversity of *Salmonella* strains recovered from stool samples of eight patients, with up to 20 isolates analysed per sample.

A total of 156 *Salmonella enterica* isolates were recovered. All isolates from individual patients displayed consistent serovars and sequence types. Despite the serotype consistency, microevolution was observed in *S.* Java ST149 and *S.* Java ST43, with SNP analyses revealing higher diversity (13 and 5 SNP differences, respectively) compared to the clonal populations of other serovars. Phylogenetic analysis of *S.* Java ST149 isolates from Patient 2 revealed distinct branching, driven by mutations in genes such as *secY* and *cnoX*, while *S.* Java ST43 isolates from Patient 1 displayed multiple clades with notable SNPs affecting transcriptional regulators.

Genome structure (GS) analyses using hybrid assemblies identified uniform GS1.0 across all isolates. Antimicrobial resistance (AMR) profiling revealed the presence of multidrug efflux pump genes (*mdsA* and *mdsB*) in all isolates. However, *S.* Typhimurium isolates from Patient 4 exhibited additional AMR genes, including *sul2*, *aph(3’’)-Ib*, and *blaTEM-1*, associated with an 8.7 kb resistance region. A single isolate from Patient 4 lacked these additional genes due to the deletion of a ∼19 kb genomic region, highlighting structural variation as a driver of phenotypic differences.

These findings emphasise the genetic diversity of *Salmonella* within hosts, particularly in serovars such as *S.* Java, and underscore the limitations of single-colony sequencing in capturing this complexity. The study highlights the utility of hybrid sequencing strategies for comprehensive analysis of genome variation, offering valuable insights into transmission dynamics, antimicrobial resistance, and evolutionary processes in *Salmonella*.

**Impact statement:** This study provides a transformative perspective on *Salmonella* genomics, uncovering the significant within-host diversity of *Salmonella* populations through the analysis of multiple single-colony isolates. By leveraging hybrid sequencing technologies, we reveal critical insights into the microevolution of *Salmonella* during infection, capturing genome-wide variation at both single nucleotide and chromosomal scales. Our findings highlight the inherent limitations of traditional single-colony sequencing in detecting within-host diversity, particularly in genetically diverse serovars such as *S. Java* ST149.

The identification of antimicrobial resistance (AMR) determinants and structural genomic variation demonstrates the dynamic nature of *Salmonella* genomes, offering crucial implications for clinical management and public health surveillance. Importantly, we show that within-host diversity may influence epidemiological interpretations, particularly in outbreak investigations and source attribution, where genetic differences between isolates can mask deeper transmission links.

**Data Summary:** All relevant supporting data are available in the accompanying supplementary data files. The online version of this article contains four supplementary table.

All *Salmonella* isolate genome sequences are available in the National Centre for Biotechnology Information (NCBI), Sequence Read Archive (SRA) under the Bioproject accession numbers PRJNA1230128. SRA accession numbers and associated metadata for isolate genomes are included in the supplementary table S2-S4.

## Introduction

*Salmonella* is a significant public health pathogen, causing a spectrum of diseases ranging from gastroenteritis to enteric fever and invasive non-typhoidal salmonellosis (iNTS) [1]. Enteric fever is caused by the *Salmonella enterica* subsp. *enterica* serovars Typhi (*S*. Typhi) and *S*. Paratyphi A, B, and C [2, 3], while non-typhoidal invasive diseases are often associated with *S.* Choleraesuis, *S*. Dublin, *S*. Panama, *S*. Virchow, and *S*. Typhimurium ST313 [4-6]. In contrast, gastroenteritis can be caused by a wide range of non-typhoidal *Salmonella* (NTS), not limited to *Salmonella enterica* subsp. *enterica* [7]. Phylogenetic analysis of the *Salmonella* genus identifies three species (*S. bongori*, *S. arizonae*, and *S. enterica*), with *S. enterica* comprising ten subspecies: *enterica*, *salamae*, *arizonae*, *diarizonae*, *houtenae*, *indica*, *londinensis*, *brasiliensis*, *hibernicus*, *essexiensis*, and *reptilium* [7].

Symptoms of diarrhoea, mild fever, and stomach cramps are characteristic of gastroenteritis caused by *Salmonella,* termed salmonellosis. Some patients also experience nausea, vomiting, headaches and muscle pains [8]. In the United Kingdom (UK), most cases are described as mild with symptoms lasting 4-7 days [9]. Individuals with prolonged symptoms, as well as at-risk groups such as the elderly, young children, and immunocompromised individuals, may require medical intervention to manage the infection, particularly due to the risks posed by invasive strains capable of causing bacteraemia [10].

Infectious dose estimates vary according to serovar, health, age, and the vehicle of transmission. Generally, the infectious dose is considered to be between 10^3^ and 10^5^ viable bacterial cells [11, 12]. Common sources of *Salmonella* include poultry, meat, eggs and egg products, dairy products, processed foods, and contaminated water [13, 14]. Other sources of note are pets such as reptiles, amphibians and dogs as well pet food [13, 15].

In the UK, *Salmonella*-induced gastroenteritis causes an estimated 38,000 community cases annually [16]. Each year, an average of 8,000 cases of salmonellosis are reported in England and Wales through general practitioners (GPs) and hospital inpatients [17, 18]. In 2022, the UK Health Security Agency (UKHSA) documented 11 *Salmonella* outbreaks in England, affecting 591 people and resulting in 3 deaths, with food sources identified in 10 of the outbreaks [18]. During the same time period, the European Union reported 1,014 *Salmonella* outbreaks, with 65,208 cases and 81 deaths. Among these, 200 cases were classified as having strong evidence of source attribution, with 151 outbreaks linked to specific food vehicles [19]. Tracing outbreaks to their source is essential for controlling the spread of infection; however, this task is hindered by the genetic diversity of *Salmonella* and the intricate nature of global food supply chains [20, 21].

High-throughput short-read whole genome sequencing (WGS) technologies have enabled a transition from biochemical based typing methods to analysis of DNA sequences [22]. The reliability and high resolution of DNA sequence-based analysis have prompted public health agencies around the world to adopt DNA sequencing as their gold standard method for surveillance and outbreak investigations [22-28].

The standard protocols for WGS analysis involve selecting a single colony from a culture plate as input material for DNA extraction and sequencing [28-30]. While this approach provides an accurate representation of a single *Salmonella* genome, within-patient *Salmonella* diversity cannot be assessed. This is particularly important in cases where multiple *Salmonella* strains may coexist within a patient or an outbreak. This convention creates a gap in our knowledge of *Salmonella* diversity within a patient.

Recent studies have identified genome-level diversity within a single host infection for various human pathogens, including *Burkholderia dolosa* [31]*, Campylobacter* [32], *Clostridium difficile* [33], *Helicobacter pylori* [34], *Mycobacterium tuberculosis* [35], *Staphylococcus aureus* [36, 37], and *Streptococcus pneumoniae* [38]. If a patient is infected with multiple strains, sequence types, or a population containing significant single nucleotide polymorphisms (SNPs), our ability to effectively conduct surveillance and accurately reconstruct transmission chains from a single colony is compromised.

Among the numerous sequence types (STs) of *Salmonella enterica*, certain lineages exhibit marked differences in genetic diversity, reflecting their evolutionary histories, ecological niches, and host ranges [39]. For instance, *S.* Enteritidis ST11 is highly clonal and globally distributed, driven by its association with poultry and widespread dissemination through the food industry [40-42]. Similarly, *S.* Java ST43, often linked to aquaculture and livestock, is relatively clonal due to selective pressures such as antimicrobial use [43]. In contrast, *S.* Typhimurium ST34 demonstrates significant genetic diversity, fuelled by its broad host range, environmental adaptability, and frequent acquisition of mobile genetic elements, including antimicrobial resistance genes [44, 45]. Lesser-known sequence types, such as *S.* Java ST149 and *S.* Anatum ST5197, remain poorly characterised, potentially representing more diverse groups due to less-defined ecological niches. Beyond *S. enterica* subsp. *enterica*, subspecies such as *S. enterica* subsp. *salamae* also display substantial genetic diversity, underscoring the potential for other subspecies to rival the diversity seen in *S. enterica* subsp. *enterica* [7, 46]. These variations in genetic diversity underscore the complexity of *Salmonella* populations and highlight the importance of genomic tools for accurate characterisation and surveillance.

An emerging source of variation present in bacterial genomes is genome structure. Genome structure (GS) variants occur when large genome fragments rearrange around repeat sequences such as ribosomal operons, insertion sequence (IS) elements, transposases, duplicated genes, and/or prophages [47]. Alteration of the order and/or orientation of fragments leads to many structural variants of the same genome sequence. In the case of *Salmonella*, the genome typically organises around the seven ribosomal operons. Variation in GS in *Salmonella* has been shown to affect growth phenotype and gene expression [48]. The identification of GS has been significantly enhanced by long-read sequencing platforms from Oxford Nanopore and Pacific Biosciences. Hybrid sequencing strategies now enable the assessment of genome variation at both the single nucleotide level and the chromosomal scale [48, 49].

In this study, we aim to investigate the genome-level diversity of *Salmonella* strains collected from stool samples of eight individual patients, capturing within-host genomic variation through the analysis of up to 20 isolates per sample.

## Material and Methods

### Stool collection

This study was conducted under the ethics approval of the University of East Anglia Research Ethics Committee (Ref. 2018/19-159). Human tissue (stool) research was conducted under Norwich Biorepository licence NRES number – 19/EE/0089; IRAS Project ID – 259062 approved by the UK’s Human Tissue Authority (HTA). The National Health Service (NHS) Eastern Pathology Alliance (EPA) network diagnostic laboratory in Norwich, UK, was the sole participating diagnostic laboratory. *Salmonella* spp. were initially identified from the stool specimens by the diagnostic laboratory using a rapid automated PCR-based culture-independent testing panel (Gastro Panel 2, EntericBio, Serosep United Kingdom). Once *Salmonella*-positive PCR results were confirmed, the diagnostic laboratory conducted reflexive culture according to their standard operation procedures. Eight surplus diarrhoeal stool specimens were collected from the EPA diagnostic laboratory representing eight unique anonymised patients with gastroenteritis symptoms who submitted specimens to the laboratory between March 2020 and August 2022. For this study, a 5 mL aliquot of stool was placed into a sterile specimen container, transported to the Quadram Institute Bioscience and subjected to culture-based isolation within 5 days of stool specimen submission to the EPA.

### Bacterial isolation

Stool specimens were cultured for *Salmonella* using two rounds of plating on selective media. A 10 μl aliquot of each stool specimen was directly plated to bi-plates, one half containing Xylose Lysine Deoxycholate agar (XLD) agar (Oxoid, UK) and the other half containing Brilliance*™ Salmonella* Agar (BSA; Oxoid, UK). Incubation was at 37°C for 24 hours for all steps. Up to 20 colonies expressing typical *Salmonella* morphology (colonies with black centres on XLD and purple colonies on BSA) from either agar were transferred to MacConkey agar (Oxoid, UK) for purification. As a final purification step, putative *Salmonella* isolates were cultured onto Tryptic Soy Agar (TSA) plates (Oxoid, UK). Isolates were preserved in Brucella broth supplemented with 17.5% glycerol.

### DNA extraction

DNA extraction was carried out using the Fire Monkey HMW DNA extraction kit (RevoluGen, Hadfield, UK) in 96-well format utilising the positive air pressure on the Resolvex A200 robotic platform (Tecan) using an adapted method: STET1 (8% sucrose, 50 mM Tris-HCl, 50 mM EDTA, 5% Triton X-100) buffer was utilised with 30mg/ml lysozyme during the lysis step. The DNA was quantified using Quant-IT broad range (Thermo Scientific, Paisley, UK) with a GloMax® Discover Microplate Reader (Promega, Southampton, UK). A subset of samples was analysed with FemtoPulse (Agilent Technologies, Wokingham, UK) to confirm presence of high molecular weight DNA.

### Sequencing

A miniaturised library preparation method was employed for short-read sequencing using Illumina technology. DNA for sequencing was prepared using the Illumina DNA Prep (Illumina Ltd, Cambridge, UK) as previously described [50]. Paired-end 150bp sequencing was carried out on a NextSeq 550 platform (Illumina Ltd, Cambridge, UK). For long read sequencing isolate DNA was sequenced in batches of 48 on a MinION using R9.4.1 flow cells (FLO-MIN106) in conjunction with Ligation Sequencing Kit (SQK-LSK109) and Native Barcoding Expansion 96 kit (EXP-NBD196) (Oxford Nanopore Technologies, Oxford, UK). Flow cells were run for 72 hours. Raw sequencing data was collected using ONT MinKNOW software (v4.0.5) and subjected to local base-calling, de-multiplexing and barcode trimming using ONT Guppy (v5.0.11).

### Hybrid genome assembly

Short reads were quality controlled using fastp (v0.19.5) and visualised using MultiQC (v1.11) [51, 52]. Long-read sequences were filtered for high-quality using Filtlong (v0.2.0) [53] with Illumina short reads sets used as external reference with Min. length = 1000, Min. mean quality = 10, and Trim non-k-mer-matching activated. Hybrid assembly was run using Unicycler (v0.4.8.0) [54]. The NCBI prokaryotic genome annotation pipeline was used to annotate genomes with *Salmonella* set as the genus [55].

### In silico subtyping, AMR predictions and variation analysis

Sequence analysis was performed on the open-source Galaxy platform (REF). *In silico* typing was carried out using SeqSero2 (Galaxy Version 1.2.1) [56] and MLST (Galaxy Version 2.16.1) [57]. Antimicrobial resistance genes were predicted using abriTAMR (Galaxy Version 1.0.14) (DOI <u>10.5281/zenodo.7370627</u>). The order and orientation of each chromosome was analysed using *socru* (Galaxy Version 2.2.4) [47] or by manual alignment and visualisation in Artemis Comparison Tool (v18.0.2) [58]. Single nucleotide polymorphism analysis was carried out using snippy4/snippy-core (Galaxy Version 4.4.3, https://github.com/tseemann/snippy). IQ-tree (Galaxy Version 2.1.2) [59, 60] as used for tree construction. For each patient one hybrid assembly was selected to be a reference based on quality metrics including read coverage, complete genome structure solved using *socru* [47] and low contamination score obtained from CheckM (Galaxy Version 1.0.11) [61]. Paired short read sequence data were used to align to the reference per patient. Maximum likelihood trees were visualised with iTOL (https://itol.embl.de/).

### Evaluating cases of Salmonella enterica subsp. salamae in England and Wales

To evaluate the incidence of *Salmonella enterica* subsp. *salamae* in England and Wales a dataset consisting of 356 strains was identified using filtering parameters with Enterobase [62]. *Salmonella* entries were filtered to include PHE (UKHSA) and the GBRU (Gastrointestinal Bacteria Reference Unit) as Lab Contact and II as Subspecies. Entries with Multi Locus Sequence typing (MLST) information were used to from an MSTree2 embedded within Enterobase. Entries with assembly data were downloaded and analysed using abriTAMR (Galaxy Version 1.0.1).

## Results

### Patient characterisation

Stool specimens from 8 patients were PCR-confirmed salmonellosis cases presenting with diarrhoea. Five cases submitted specimens to general practitioners, while the remaining three were submitted at Norfolk and Norwich University Hospital. Notably, two patients reported a recent travel history. The patients ranged from 2 to 77 years of age, and 7 out of 8 cases identified as female (Table S1).

### Salmonella genomic subtyping within patients

A total of 156 *Salmonella* isolates were recovered. The number of isolates per patient stool specimen ranged from 18 to 20 isolates. All isolates recovered were *Salmonella enterica* subsp. *enterica,* except for one patient who harboured *Salmonella enterica* subsp. *salamae.* SeqSero2 and MLST were used to obtain a serotype prediction and sequence type per isolate. For all isolates from each patient, a single serotype and sequence type was observed including two cases of *Salmonella enterica* serovar Paratyphi B variant Java and two cases of *Salmonella enterica* serovar Enteritidis (Table 1).

**Table 1.** Description of *Salmonella* identified in stool samples of eight gastroenteritis patients in Norfolk, UK.

| Patient ID | *Salmonella enterica* subsp. | Sequence Type (ST) | No. of isolates per patient |
| --- | --- | --- | --- |
| Patient 1 | *Salmonella enterica* subsp. *enterica* serovar Paratyphi B variant Java | 43 | 20 |
| Patient 2 | *Salmonella enterica* subsp. *enterica* serovar Paratyphi B variant Java | 149 | 19 |
| Patient 3 | *Salmonella enterica* subsp. *enterica* serovar Infantis | 32 | 18 |
| Patient 4 | *Salmonella enterica* subsp. *enterica* serovar Typhimurium | 34 | 20 |
| Patient 5 | *Salmonella enterica* subsp. *enterica* serovar Enteritidis | 11 | 20 |
| Patient 6 | *Salmonella enterica* subsp. *salamae* | 9581 | 20 |
| Patient 7 | *Salmonella enterica* subsp. *enterica* serovar Anatum | 5197 | 20 |
| Patient 8 | *Salmonella enterica* subsp. *enterica* serovar Enteritidis | 11 | 19 |

### Maintenance of genome structure

To identify if GS was uniform across the isolates obtained from each patient the GS was solved by running *socru* on hybrid genome assemblies. Across all eight patients the GS of isolates were uniform with no differences observed. The GS observed for all isolates was GS1.0 [47].

### AMR determinants

All isolate genomes were examined for antimicrobial resistance (AMR) determinants using complete hybrid genome assemblies. Regardless of serovar, all genomes exhibited the presence of the *mdsA* and *mdsB* genes, which encode a multidrug resistance efflux pump [63]. For Patient 4, 19 out of 20 *S.* Typhimurium isolates displayed a broader AMR determinant profile, carrying additional resistance determinants of *sul2* (sulfonamide resistance), *aph(3’’)-Ib* and *aph(6)-Id* (aminoglycoside resistance), and *blaTEM-1* (beta-lactam resistance) genes. The 8.7kb region carrying the AMR genes *sul2, aph(3’’)-Ib,aph(6)-Id, blaTEM-1* was flanked by and contained IS15DIV transposase insertion sequences. One *S.* Typhimurium isolate out of the 20 (Isolate 10, Patient 4) contained only *mdsA* and *mdsB.* Inspection of the genome revealed a single insertion sequence at the site where *sul2, aph(3’’)-Ib,aph(6)-Id, blaTEM-1* were expected (Figure 1). Comparative genomics showed a ∼19kb region missing in this isolate’s genome which housed the four AMR genes alongside flagellar phase variation proteins *hin, fljA*, *fljB,* and a PTS transport system.

**Figure 1.**
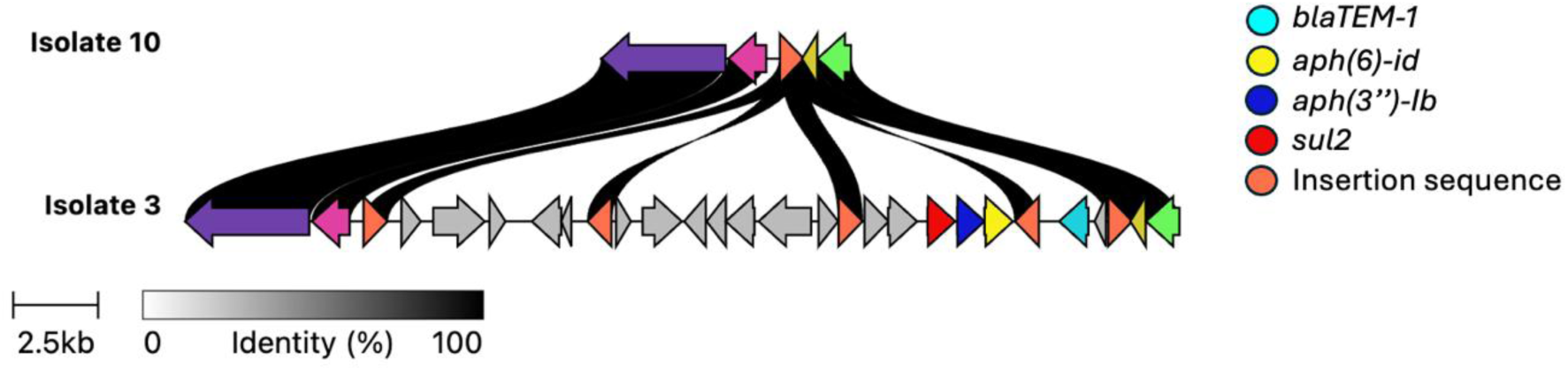
Loss of AMR-carrying transposable element in the *S*. Typhimurium genome of Patient 4. Clinker [78] schematic where isolate 3 represents the consensus sequence found in 19/20 isolates from Patient 4. Flanked by insertion sequences (in orange) several genes including four AMR genes (in red, dark blue, yellow and light blue) were absent in isolate 10. Genes are represented by arrows indicated directionality, with matching colours indicating identical gene sequence. Homology between the two isolates is represented as black bars, regions without black bars linking them are absent in isolate 10.

### SNP diversity

Based on the SNP analysis, all *Salmonella* isolates recovered from Patients 3-8 were considered clonal with a maximum SNP difference of one. *Salmonella* Java recovered from Patients 1 and 2 were more diverse, with a maximum SNP difference of five and thirteen, respectively (Table 2). Across the dataset, a total of 38 SNPs were observed; 26 were missense variants, three caused truncation by gaining a STOP codon, two were in non-coding DNA, and seven were synonymous (Table 3).

**Table 2.** Number of SNPs identified between isolates within each stool sample of eight gastroenteritis patients in Norfolk, UK.

| Isolate ID |  |  |  |  |  |  |  |  |  |  |  |  |  |  |  |  |  |  |  |  |  |  |
| --- | --- | --- | --- | --- | --- | --- | --- | --- | --- | --- | --- | --- | --- | --- | --- | --- | --- | --- | --- | --- | --- | --- |
| Patient ID | *Salmonella*<br>serovar | Sequence type<br>(ST) | 1 | 2 | 3 | 4 | 5 | 6 | 7 | 8 | 9 | 10 | 11 | 12 | 13 | 14 | 15 | 16 | 17 | 18 | 19 | 20 |
| Patient 1 | Java | 43 | 0 | 0 | 0 | 3 | 3 | 1 | 0 | R | 2 | 1 | 2 | 5 | 2 | 1 | 0 | 3 | 0 | 2 | 3 | 3 |
| Patient 2 | Java | 149 | 9 | 0 | 9 | 9 | 9 | 9 | 9 | 9 | 9 | 9 | R | 9 | 9 | 0 | 9 | 0 | n/a | 0 | 13 | 0 |
| Patient 3 | Infantis | 32 | 1 | 0 | 0 | 1 | 0 | 0 | 0 | n/a | 0 | 1 | 0 | R | 0 | 1 | 0 | 0 | 0 | 0 | n/a | 0 |
| Patient 4 | Typhimurium | 34 | 0 | 0 | R | 0 | 0 | 0 | 0 | 0 | 0 | 0 | 0 | 0 | 0 | 0 | 0 | 0 | 0 | 0 | 0 | 0 |
| Patient 5 | Enteritidis | 11 | 0 | R | 1 | 0 | 1 | 1 | 0 | 1 | 0 | 0 | 0 | 0 | 0 | 0 | 0 | 0 | 0 | 0 | 0 | 0 |
| Patient 6 | *salamae* | 9581 | 0 | 0 | 0 | 0 | 0 | 0 | 0 | 0 | 0 | 0 | 0 | 0 | 0 | 0 | 0 | 0 | 0 | 0 | 0 | R |
| Patient 7 | Anatum | 5197 | 0 | 0 | 0 | 0 | 0 | 1 | 0 | 0 | 0 | 0 | 0 | 0 | R | 0 | 0 | 0 | 0 | 1 | 0 | 0 |
| Patient 8 | Enteritidis | 11 | 0 | 0 | 0 | 0 | 0 | 0 | R | 0 | 0 | 0 | 0 | 0 | n/a | 0 | 0 | 0 | 0 | 0 | 0 | 1 |

**R** = reference strain, **n/a** = no sequence data for isolate

**Table 3.**
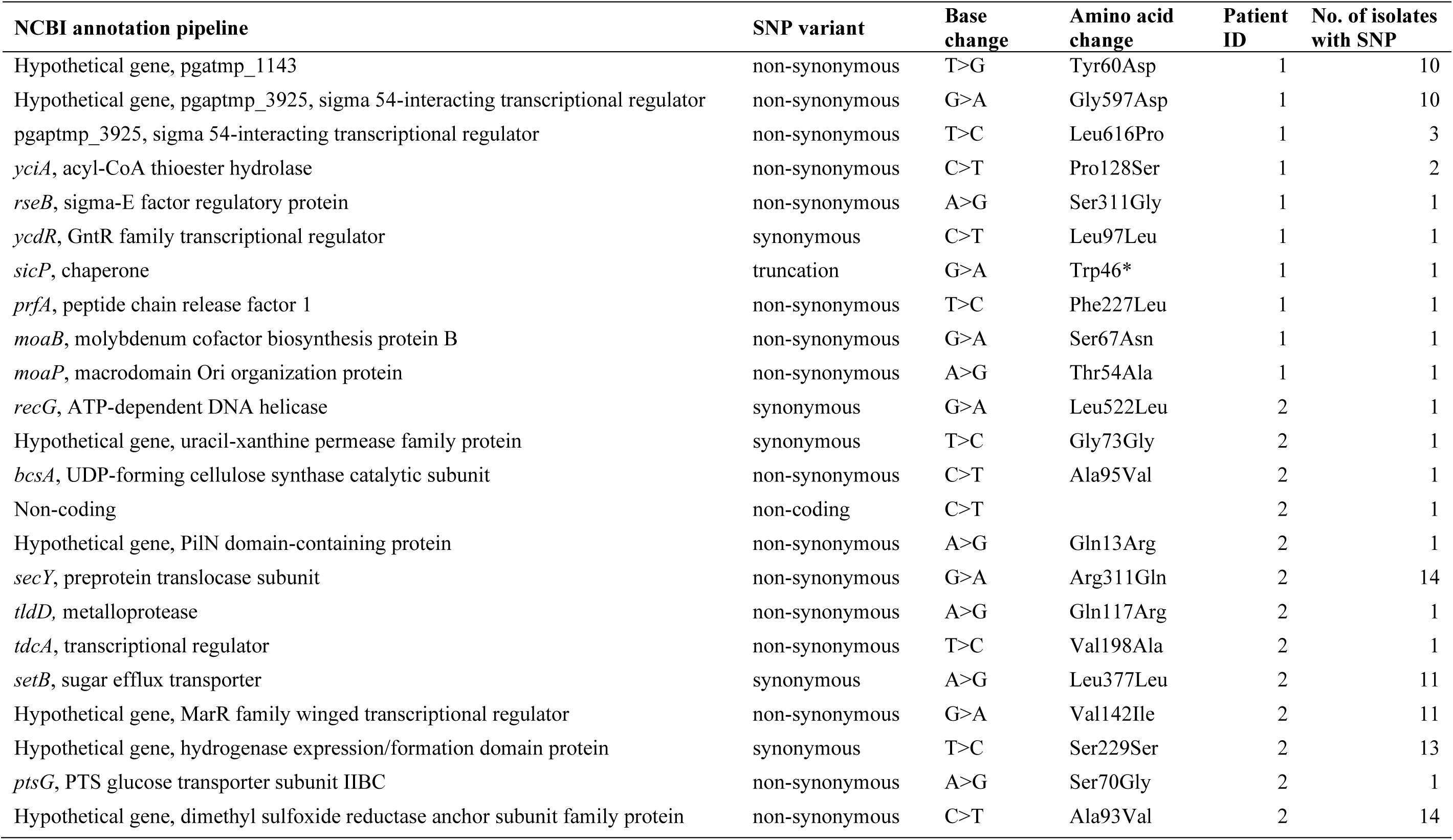
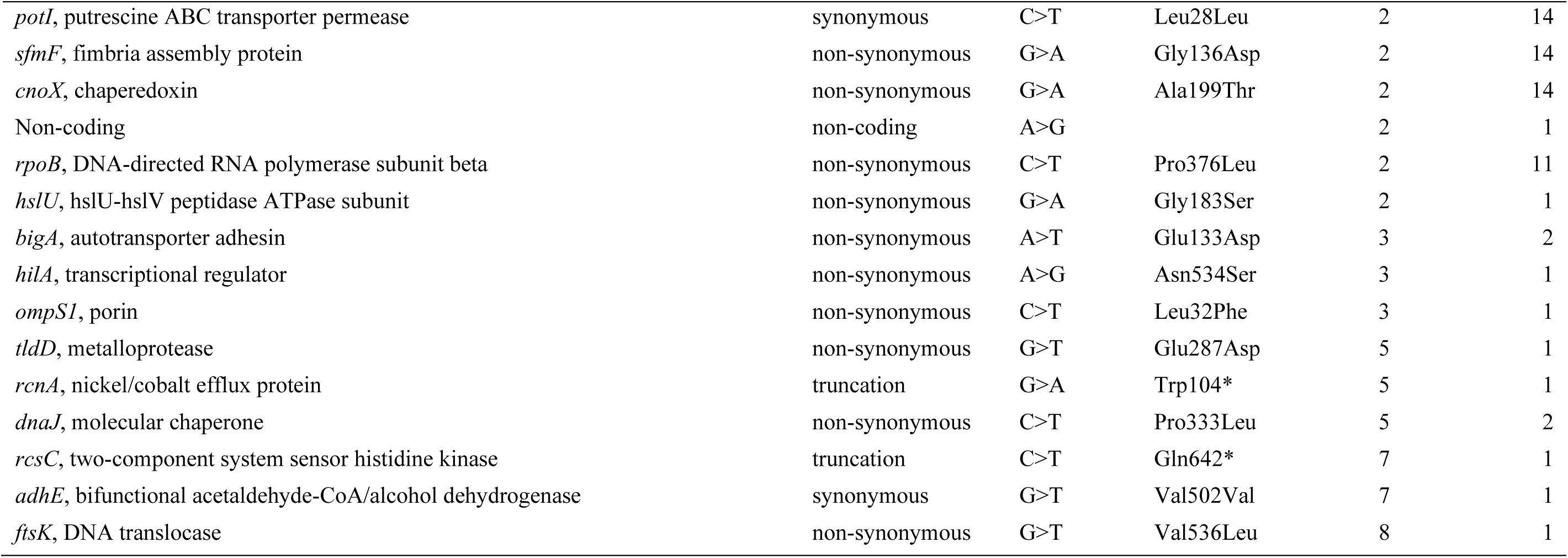
Summary of SNPs identified in isolates from eight patients with salmonellosis in Norwich, UK, 2020-2022.

The most diverse isolates were the *S*. Java ST149 isolates from Patient 2. The core genome phylogenetic tree revealed a process of microevolution occurring (Figure 2). A deep branch separated a basal group of six isolates from the remaining thirteen. The six SNPs responsible for forming this branch were in *secY*, *potI(ydcV)*, *sfmF, cnoX*, a hypothetical gene encoding a dimethyl sulfoxide reductase anchor subunit family protein, and a hypothetical gene encoding a hydrogenase expression/formation domain-containing protein. Beyond the basal branch, three separate branches were present. Isolate 19 formed its own branch with a seven SNP distance, Isolate 13 formed its own branch with a three SNP distance, and a group of eleven isolates formed a clade at a distance of three SNPs.

**Figure 2.**
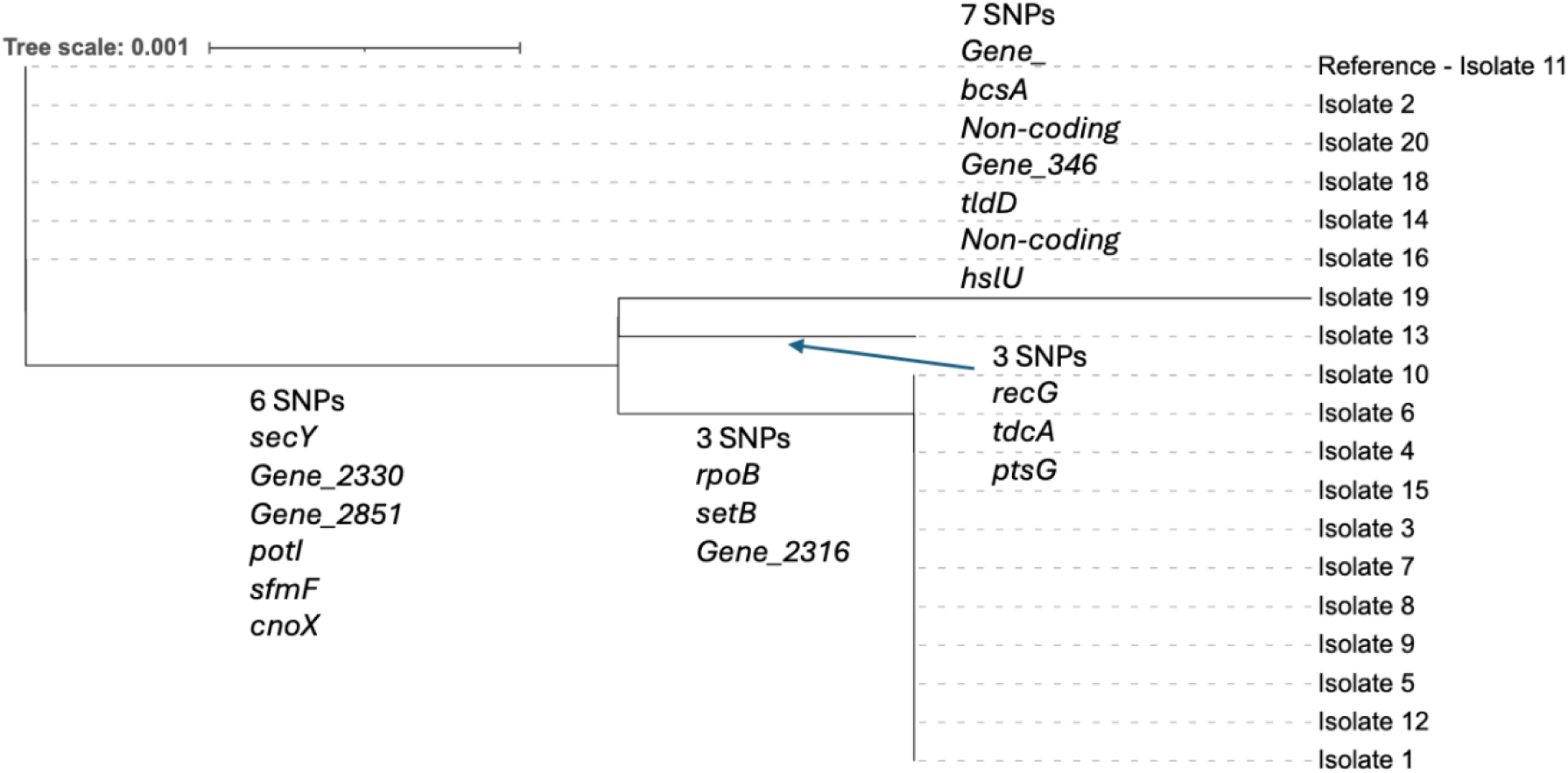
Within-patient variation in Patient 2 (*S.* Java). Core genome maximum likelihood tree for nineteen *S.* Java isolates (Patient 2). Tree overlaid with SNPs responsible for each branch.

Three basal branches were observed in the core SNP tree for the isolates from Patient 1’s *S.* Java ST43 infection: Isolate 14 appeared on its own branch, Isolate 6 and Isolate 11 formed another branch, and a clade of ten isolates appeared on a third branch. This clade of ten isolates was further diversified with four more branches. Isolate 4, Isolate 5, and Isolate 19 carried a double mutation in *Gene_3925* (Figure 3). The gene was predicted to act as a transcriptional regulator for a PTS sugar transporter and was located upstream of the predicted PTS system subunits IIA, IIB, IIC, and IID. Both SNPs in *Gene_3925* resulted in missense variants within the PTS EIIA mannose/sorbose-specific type-4 domain of the protein, a component of the enzyme II (EII) complex.

**Figure 3.**
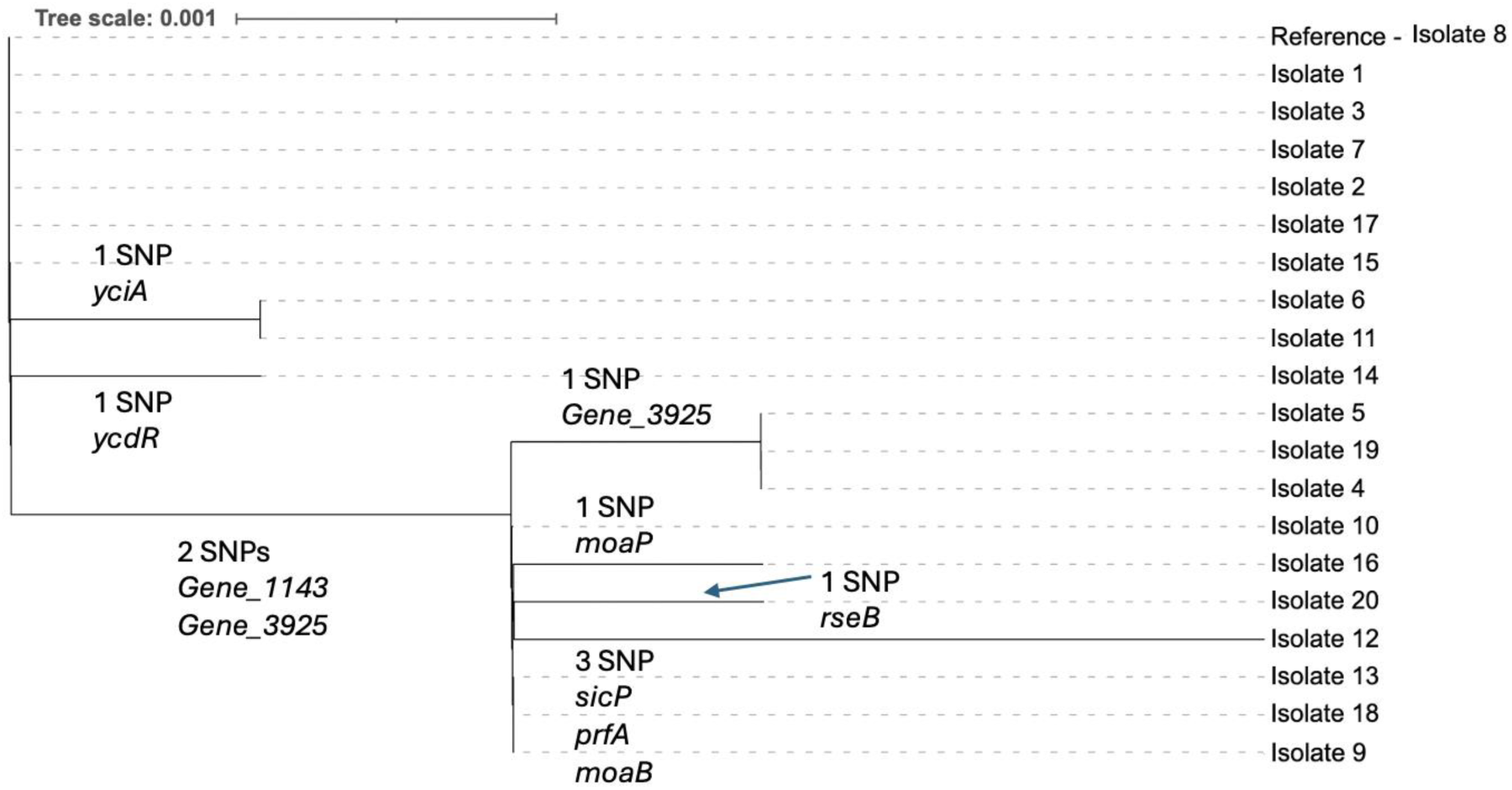
Within-patient variation in Patient 1 (*S.* Java). Core genome maximum likelihood tree for twenty *S.* Java isolates (Patient 1). Tree overlaid with SNPs responsible for each branch.

Additional mutations that appeared in multiple isolates within a single patient group included a missense variant in *bigA*, found in two *S.* Infantis isolates from Patient 3, and a missense variant in *dnaJ*, observed in two *S.* Enteritidis isolates from Patient 5. The gene *bigA* encodes a large putative surface-exposed virulence protein associated with virulence in some intracellular pathogens [64]. The gene *dnaJ* is located within the *Salmonella* pathogenicity island (SPI) II. Loss-of-function mutations in the *dnaJ* gene have been shown to increase heat resistance in *S*. Typhimurium [65]. A mutation in the metalloprotease *tldD* was identified in a single isolate from Patient 2 (Java) and Patient 5 (Enteritidis). Both mutations resulted in missense variants affecting different domains within the TldD protein. Three mutations resulting in truncated proteins were identified: in a single isolate from Patient 1 (Java), SicP was truncated from 130 amino acids to 43 amino acids; in a single isolate from Patient 5 (Enteritidis), RcnA was truncated from 284 amino acids to 104 amino acids; and in a single isolate from Patient 7 (Anatum), RcsC was truncated from 948 amino acids to 642 amino acids.

### Rare case of gastroenteritis caused by Salmonella enterica subsp. salamae in the UK

There are 356 subspecies II (*salamae*) entries in Enterobase linked to England and Wales deposited by PHE (UKHSA) and the GBRU with MLST information. Of the 356 entries, 123 different sequence types were observed. The most commonly observed STs were ST53 and ST8955, with 30 and 28 entries, respectively (Figure 4). Patient 1’s isolates were ST9581 which had a single entry. Based on MLST analysis ST9581 was most closely related to ST2307, a serovar seen 15 times between 2015 and 2024. Analysis of the AMR profiles of the 285/356 genome assemblies available from Enterobase revealed low levels of AMR determinants among subspecies II linked to gastroenteritis in the UK. The *fosA7.4* gene was observed in 27 isolates, *fosA7.2* in one isolate, and *tet(B)* in one isolate. The efflux pairing *mdsA* and *mdsB* present in all of Patient 6’s *S. enterica* subsp. *salamae* isolates was observed in 200 isolates, leaving 85 isolates with no recognisable AMR determinants.

**Figure 4.**
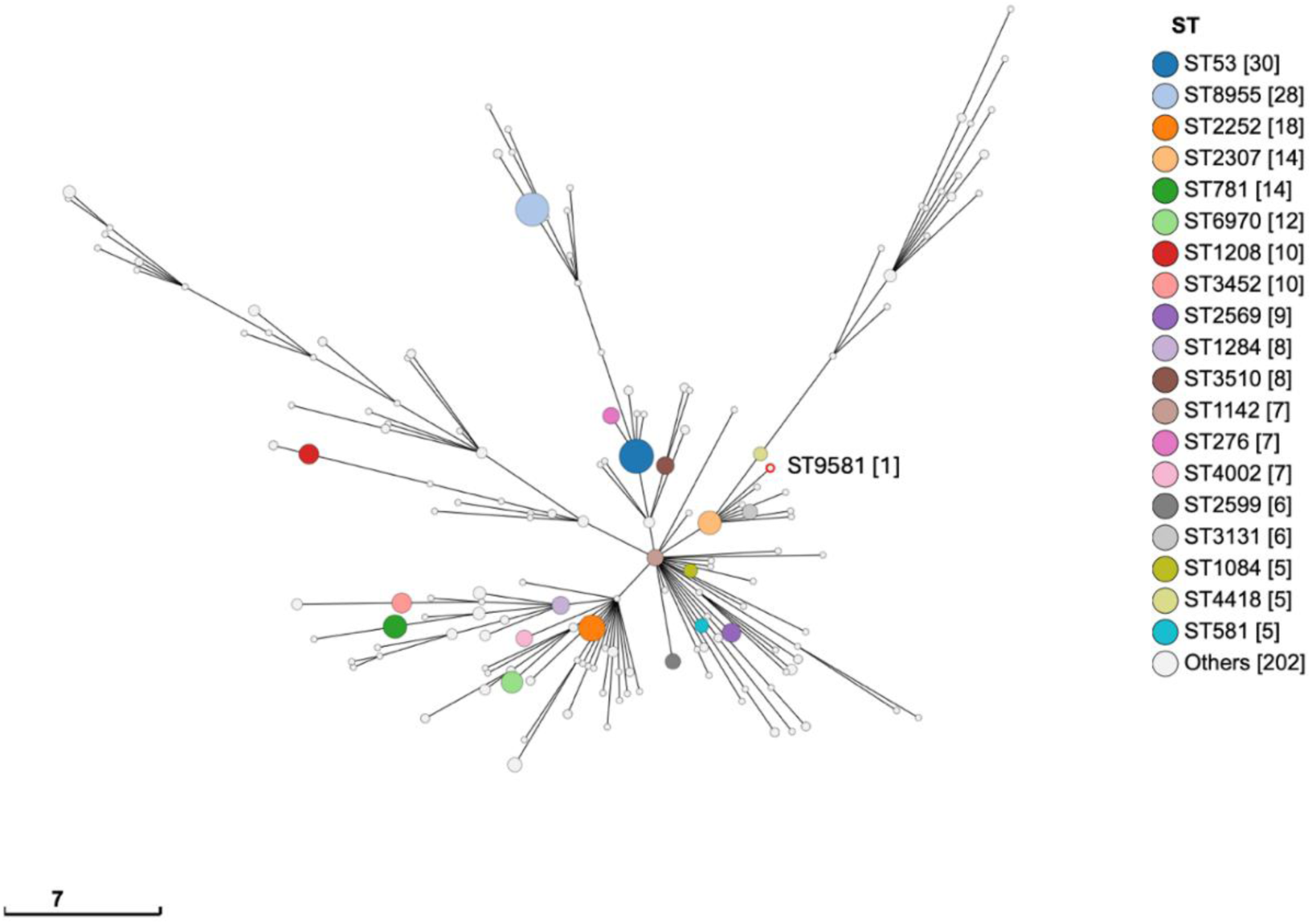
Phylogenetic relationships within *S. enterica* subsp. *salamae* from England & Wales. Achtman 7 Gene MLST MSTree2 of Subspecies II entries submitted to Enterobase by GBRU/PHE, 1956-2024. Patient 1’s isolate outlined in red and labelled, ST[Occurrence].

## Discussion

To our knowledge this is the first study analysing multiple isolates of *Salmonella* identified from individual patients’ stool samples who were suffering from gastroenteritis. This study analysed genomic diversity of *Salmonella* within eight patient’s stool samples considering subspecies, sequence type, serotype, genome structure, AMR profile, and SNPs. No mixed *Salmonella* species infections were observed, with all isolates from individual infections being identified as the same subspecies, serovar, and sequence type. This aligns with the existing literature on *Salmonella*, where reports of multiple-serovar infections are exceptionally rare and seldom documented [66, 67].

The genome structure of all genomes screened within a stool sample was uniform, and identified as GS1.0 and is in alignment with GS1.0 being the most commonly observed genome structure across the *Salmonella* genus [47]. Structural deviations from GS1.0 are overrepresented in *S. Typhi* and have been linked to persistence within the human host [47, 68]. Recent analysis of *S*. Agona isolates from UK infections found GS1.0 to be the most dominant structure [49]. The study included isolate sequencing for acute and persistent infections within individual patients linking alterations in genome structure away from GS1.0 to early, convalescent carriage (3 weeks– 3 months). As an emerging source of genome level variation, the impact of genome structure on infection severity, progression and host persistence remains unknown. To our knowledge, all patients in this study suffered from acute disease only.

Our study reveals that SNPs were detected in *Salmonella* isolates from six out of eight patients during short-term passage through the gastrointestinal tract. The isolates recovered from Patient 4 (*S*. Typhimurium) and Patient 6 (*S. enterica* subsp. *salamae*) had no SNP differences in a core alignment. The most diverse set of isolates were from Patient 2’s *S*. Java ST149 infection. Two clades appeared to be present in this population separated by nine SNPs, with isolate Isolate-19 having a thirteen SNP distance from the reference. For outbreak investigations involving *Salmonella*, UKHSA use single linkage clustering at 250, 100, 50, 25, 10, 5 and 0 SNPs, forming a SNP address [22, 69]. Thresholds of 0 and 5 SNPs are deemed to support a meaningful relatedness between isolates and their point of origin. At a threshold of 10 SNPs, the relationship between isolates is less clear, although it is noted that deeper epidemiological links may be uncovered by analysis at this level, providing important clues during an outbreak investigation [69]. A key takeaway from this study is that the genome-level diversity present among Patient 2’s *S. Java* ST149 may not be accurately observed when sequencing a single isolate.

Patients 1’s *S*. Java ST43 infection formed two clades separated by two SNPs with a maximum distance from the reference of five SNPs. Three isolates carried two non-synonymous SNPs in *Gene_3925*, a sigma 54-interacting transcriptional regulator similar to the *dgaR* gene. The *dgaR* gene has previously been described as an RpoN-dependent activator of the *dgaABCDEF* operon, a mannose family phosphotransferase system (PTS) enabling the catabolism of d-glucosaminate to pyruvate plus glyceraldehyde-3-phosphate [70]. All PTS permeases consist of three domains, named EIIA, EIIB, and EIIC, whereas mannose family PTS permeases include a fourth domain, referred to as EIID [71, 72]. EII complexes are linked to the cell membrane and are specific to a particular sugar or a group of structurally similar sugars [70]. In the case of the operon led by *Gene_3925*, EIIA is a hypothetical PTS sugar transporter subunit, EIIB is annotated as a sorbose-specific EIIB component, EIIC is annotated as a N-acetylgalactosamine permease IIC, and EIID is annotated as a PTS system mannose-specific EIID component. The dual mutation observed to *Gene_3925* is suggestive of adaptive pressure on this pathway within S*. Java* ST43 during passage through the human digestive system.

Observation of different AMR profiles for isolates from the same patient were rare. For seven of the eight patient stool samples, the *Salmonella* isolates exhibited limited AMR determinants, carrying only the efflux pump genes *mdsA* and *mdsB*. In contrast, Patient 4’s *S*. Typhimurium infection showed a notable difference in its AMR profile, with one isolate predicted to be sensitive, while the consensus profile included *sul2*, *aph(3’’)-Ib*, *aph(6)-Id*, and *blaTEM-1*. Long-read sequencing enabled the assembly of complete circular genomes, allowing for detailed inspection of the genome sequences of these isolates, including location of repetitive elements and AMR determinants. Analysis revealed that these four AMR genes were located within a genomic region flanked by and containing five IS15DIV insertion sequences (Figure 1). IS15DIV has been associated with co-occurrence of resistance markers in *E. coli* [73]. A BLAST search of the region revealed 100% homology to a plasmid from an Enterohemorrhagic *Escherichia coli* O111:H8 strain recovered from a large outbreak in Japan associated with the consumption of raw beef [74]. This transposable element has also been identified within a chromosomal drug resistance region in an ESBL-producing *Salmonella Typhi* isolate (814995) from a UK Patient with enteric fever symptoms, linked to travel history in Karachi, Pakistan, in September 2019 [75]. We hypothesise that the sensitive *S*. Typhimurium ST34 isolate lost the four AMR genes in a recombination event leaving a single insertion sequence at this location. Observing different *Salmonella* AMR profiles within a single infection demonstrates that analysing a single colony may not accurately represent the *Salmonella* population responsible for the infection and impedes investigative conclusions. If the sensitive colony was selected as representative of the population, then valuable information about the resistance profile would have been missed. While sequencing multiple isolates individually increases the investigation power it remains unclear how many isolates would be needed to fully capture the genetic diversity within a *Salmonella* population during an infection. For *Campylobacter* it has been suggested that up to 80 isolates would be needed to capture 95% of core non-recombinant SNPs [32].

A significant cost is associated with sequencing multiple isolates per patient sample. Additional costs of labour, isolating the multiple colonies, DNA extraction, DNA sequencing, and data storage make a multiple colony approach unfeasible for any large-scale diagnostic pipelines. Alternative sampling strategies such as sweep sequencing or pool-seq where multiply isolates are DNA extracted and sequenced as one sample have been proposed as an alternative approach capable of capturing information at the population level [36, 76, 77]. These alternative approaches would alleviate the detection limitations of using a single colony; however, both methods have their own constraints. Sweep sequencing can result in the loss of minor alleles, leading to a biased representation of genetic diversity. Similarly, pool-seq, while capable of capturing a broader representation of population diversity, can introduce biases during pooling due to unequal DNA contributions from individual isolates, and low-frequency variants may remain undetected unless sequencing depth is sufficiently high. Furthermore, neither method provides insights into the genomic context of individual isolates, limiting their ability to resolve detailed genome structure and link SNPs and/or AMR profiles to individual isolates.

## Conclusion

Our sequencing of multiple *Salmonella* isolates from patient stool samples demonstrated varying levels of genome diversity within the population in different *Salmonella* serovars and sequence types involved in gastroenteritis cases. This is informative to epidemiological investigations involving sequence types displaying more genetic diversity (*e.g. S*. Java). Sequencing strategies that enable analysis of many isolates from a single patient’s sample can provide a higher resolution of genetic information, capturing the full spectrum of genetic diversity within the bacterial population. This approach can identify subtle variations and rare mutations that might otherwise go undetected. Over time, such detailed analysis could help uncover common genes under selective pressure during the infection process, shedding light on the mechanisms driving bacterial adaptation, persistence, and resistance.

## List of abbreviations

%: percent
°C: Degree Celsius
AMR: antimicrobial resistance
bp: base pair
DNA: deoxyribonucleic acid
EDTA: Ethylenediaminetetraacetic acid
EPA: Eastern Pathology Laboratory
GBRU: Gastrointestinal Bacteria Reference Unit
GP: General practitioner
GS: Genome structure
HCl: Hydrogen chloride
HMW: High molecular weight
HTA: Health Technology Assessment
IS: Insertion seqeunce
iTOL: Interactive Tree Of Life
kb: kilobase
mg: milligram
min.: minimum
ml: millilitre
mM: millimeter
MLST: Multilocus sequence typing
PCR: polymerase chain reaction
PHE: Public Health England
Pool-seq: 
PTS: Phosphotransferase system
ONT: Oxford Nanopore Technologies
SNP: Single Nuclotide Polymorhism
Spp.: species
ST: sequence type
TSA: Tryptic soy agar
UK: United Kingdom
UKHSA: UK Health Security Agency
v: Version
WGS: whole genome sequencing

## Declarations

### Ethical Approval

This project was approved by the Faculty of Medicine and Health Sciences Research Ethics Committee of the University of East Anglia (FMH REC reference: 201819-159HT).

### Consent for publication

Not applicable.

### Availability of data and materials

The online version of this article contains 2 supplementary table (Tables S1 and S2). All sequenced *Salmonella* isolate data are available in the National Centre for Biotechnology Information (NCBI) Sequence Read Archive under the Bioproject accession numbers PRJNA1230128. Sequence read archive (SRA) accession numbers and associated metadata can be found in the supplementary material of this study (Table S2-S4).

### Author’s contributions

G.C.L. and S.J.R designed the study conception. G.C.L. and S.J.R conceptualized the plan for the manuscript. S.J.R, drafted the manuscript. All authors contributed to edits of the manuscript. S.J.R. designed figures. S.J.R and B.D. collected and processed surplus stool samples from the NHS diagnostic laboratory. S.J.R extracted DNA and performed ONT sequencing. S.J.R carried out bioinformatic analysis and interpretation.

## Acknowledgement

We are grateful to the microbiology and administrative team at the Eastern Pathology Alliance network diagnostic laboratory in Norwich, UK for providing access to samples, laboratory managers and technicians for providing laboratory resources and support and the QIB core services sequencing team for Illumina sequence library preparation and sequencing.

