## Supplemental Tables 1-4 for "Genomic diversity of non-typhoidal *Salmonella* found in patients suffering from gastroenteritis in Norfolk, UK"

**Table S1.** Patient demographics and sample collection details for *Salmonella*-positive cases in Norwich, UK, 2020-2022

| Patient ID | EPA result | Date sample collected | Date QIB Collected | Sample origin (GP vs. Outpatient) | Age | Sex | Recent travel | Travel-region |
|---|---|---|---|---|---|---|---|---|
| 1 | Sal+ | 13/01/2020 | 14/01/2020 | GP | 31 | F | Yes | Thailand |
| 2 | Sal+ | 03/03/2020 | 03/03/2020 | GP | 62 | F | No | |
| 3 | Sal+ | 21/08/2020 | 24/08/2020 | GP | 65 | M | No | |
| 4 | Sal+ | 18/02/2022 | 21/02/2022 | Outpatient | 2 | F | No | |
| 5 | Sal+ | 20/05/2022 | 23/05/2022 | GP | 77 | F | No | |
| 6 | Sal+ | 25/06/2022 | 27/06/2022 | Outpatient | 29 | F | Yes | South Africa |
| 7 | Sal+ | 20/07/2022 | 25/07/2022 | GP | 44 | F | No | |
| 8 | Sal+ | 08/08/2022 | 10/08/2022 | Outpatient | 55 | F | na | |

**Sal+ =** Salmonella positive, **GP =** General practitioner**, EPA=** Eastern Pathology Alliance, **QIB =** Quadram Institute Bioscience**, M =** Male, **F** = Female, **na** = Not available

Table S2. Accession numbers for the Illumina reads used in study

| Accession | Sample_name | Platform | Instrument_model | Filename | Filename2 |
|---|---|---|---|---|---|
| SRR32528139 | 20EPA002NSA_1 | ILLUMINA | NextSeq 550 | 20EPA002NSA-1_R1_001.fastq.gz | 20EPA002NSA-1_R2_001.fastq.gz |
| SRR32528138 | 20EPA002NSA_2 | ILLUMINA | NextSeq 550 | 20EPA002NSA-2_R1_001.fastq.gz | 20EPA002NSA-2_R2_001.fastq.gz |
| SRR32528091 | 20EPA002NSA_3 | ILLUMINA | NextSeq 550 | 20EPA002NSA-3_R1_001.fastq.gz | 20EPA002NSA-3_R2_001.fastq.gz |
| SRR32528044 | 20EPA002NSA_4 | ILLUMINA | NextSeq 550 | 20EPA002NSA-4_R1_001.fastq.gz | 20EPA002NSA-4_R2_001.fastq.gz |
| SRR32527839 | 20EPA002NSA_5 | ILLUMINA | NextSeq 550 | 20EPA002NSA-5_R1_001.fastq.gz | 20EPA002NSA-5_R2_001.fastq.gz |
| SRR32528020 | 20EPA002NSA_6 | ILLUMINA | NextSeq 550 | 20EPA002NSA-6_R1_001.fastq.gz | 20EPA002NSA-6_R2_001.fastq.gz |
| SRR32528009 | 20EPA002NSA_7 | ILLUMINA | NextSeq 550 | 20EPA002NSA-7_R1_001.fastq.gz | 20EPA002NSA-7_R2_001.fastq.gz |
| SRR32527998 | 20EPA002NSA_8 | ILLUMINA | NextSeq 550 | 20EPA002NSA-8_R1_001.fastq.gz | 20EPA002NSA-8_R2_001.fastq.gz |
| SRR32528042 | 20EPA002NSA_9 | ILLUMINA | NextSeq 550 | 20EPA002NSA-9_R1_001.fastq.gz | 20EPA002NSA-9_R2_001.fastq.gz |
| SRR32528031 | 20EPA002NSA_10 | ILLUMINA | NextSeq 550 | 20EPA002NSA-10_R1_001.fastq.gz | 20EPA002NSA-10_R2_001.fastq.gz |
| SRR32528137 | 20EPA002NSA_11 | ILLUMINA | NextSeq 550 | 20EPA002NSA-11_R1_001.fastq.gz | 20EPA002NSA-11_R2_001.fastq.gz |
| SRR32528126 | 20EPA002NSA_12 | ILLUMINA | NextSeq 550 | 20EPA002NSA-12_R1_001.fastq.gz | 20EPA002NSA-12_R2_001.fastq.gz |
| SRR32528115 | 20EPA002NSA_13 | ILLUMINA | NextSeq 550 | 20EPA002NSA-13_R1_001.fastq.gz | 20EPA002NSA-13_R2_001.fastq.gz |
| SRR32527985 | 20EPA002NSA_14 | ILLUMINA | NextSeq 550 | 20EPA002NSA-14_R1_001.fastq.gz | 20EPA002NSA-14_R2_001.fastq.gz |
| SRR32527974 | 20EPA002NSA_15 | ILLUMINA | NextSeq 550 | 20EPA002NSA-15_R1_001.fastq.gz | 20EPA002NSA-15_R2_001.fastq.gz |
| SRR32527963 | 20EPA002NSA_16 | ILLUMINA | NextSeq 550 | 20EPA002NSA-16_R1_001.fastq.gz | 20EPA002NSA-16_R2_001.fastq.gz |
| SRR32527920 | 20EPA002NSA_17 | ILLUMINA | NextSeq 550 | 20EPA002NSA-17_R1_001.fastq.gz | 20EPA002NSA-17_R2_001.fastq.gz |
| SRR32527909 | 20EPA002NSA_18 | ILLUMINA | NextSeq 550 | 20EPA002NSA-18_R1_001.fastq.gz | 20EPA002NSA-18_R2_001.fastq.gz |
| SRR32527898 | 20EPA002NSA_19 | ILLUMINA | NextSeq 550 | 20EPA002NSA-19_R1_001.fastq.gz | 20EPA002NSA-19_R2_001.fastq.gz |
| SRR32528102 | 20EPA002NSA_20 | ILLUMINA | NextSeq 550 | 20EPA002NSA-20_R1_001.fastq.gz | 20EPA002NSA-20_R2_001.fastq.gz |
| SRR32528090 | 20EPA011NSA_1 | ILLUMINA | NextSeq 550 | 20EPA011NSA-1_R1_001.fastq.gz | 20EPA011NSA-1_R2_001.fastq.gz |
| SRR32528079 | 20EPA011NSA_2 | ILLUMINA | NextSeq 550 | 20EPA011NSA-2_R1_001.fastq.gz | 20EPA011NSA-2_R2_001.fastq.gz |
| SRR32527949 | 20EPA011NSA_3 | ILLUMINA | NextSeq 550 | 20EPA011NSA-3_R1_001.fastq.gz | 20EPA011NSA-3_R2_001.fastq.gz |
| SRR32527938 | 20EPA011NSA_4 | ILLUMINA | NextSeq 550 | 20EPA011NSA-4_R1_001.fastq.gz | 20EPA011NSA-4_R2_001.fastq.gz |
| SRR32527927 | 20EPA011NSA_5 | ILLUMINA | NextSeq 550 | 20EPA011NSA-5_R1_001.fastq.gz | 20EPA011NSA-5_R2_001.fastq.gz |
| SRR32527884 | 20EPA011NSA_6 | ILLUMINA | NextSeq 550 | 20EPA011NSA-6_R1_001.fastq.gz | 20EPA011NSA-6_R2_001.fastq.gz |

| | | | | | |
|---|---|---|---|---|---|
| SRR32527873 | 20EPA011NSA_7 | ILLUMINA | NextSeq 550 | 20EPA011NSA-7_R1_001.fastq.gz | 20EPA011NSA-7_R2_001.fastq.gz |
| SRR32527862 | 20EPA011NSA_8 | ILLUMINA | NextSeq 550 | 20EPA011NSA-8_R1_001.fastq.gz | 20EPA011NSA-8_R2_001.fastq.gz |
| SRR32528066 | 20EPA011NSA_9 | ILLUMINA | NextSeq 550 | 20EPA011NSA-9_R1_001.fastq.gz | 20EPA011NSA-9_R2_001.fastq.gz |
| SRR32528055 | 20EPA011NSA_10 | ILLUMINA | NextSeq 550 | 20EPA011NSA-10_R1_001.fastq.gz | 20EPA011NSA-10_R2_001.fastq.gz |
| SRR32527860 | 20EPA011NSA_11 | ILLUMINA | NextSeq 550 | 20EPA011NSA-11_R1_001.fastq.gz | 20EPA011NSA-11_R2_001.fastq.gz |
| SRR32527849 | 20EPA011NSA_12 | ILLUMINA | NextSeq 550 | 20EPA011NSA-12_R1_001.fastq.gz | 20EPA011NSA-12_R2_001.fastq.gz |
| SRR32527847 | 20EPA011NSA_13 | ILLUMINA | NextSeq 550 | 20EPA011NSA-13_R1_001.fastq.gz | 20EPA011NSA-13_R2_001.fastq.gz |
| SRR32527846 | 20EPA011NSA_14 | ILLUMINA | NextSeq 550 | 20EPA011NSA-14_R1_001.fastq.gz | 20EPA011NSA-14_R2_001.fastq.gz |
| SRR32527845 | 20EPA011NSA_15 | ILLUMINA | NextSeq 550 | 20EPA011NSA-15_R1_001.fastq.gz | 20EPA011NSA-15_R2_001.fastq.gz |
| SRR32527844 | 20EPA011NSA_16 | ILLUMINA | NextSeq 550 | 20EPA011NSA-16_R1_001.fastq.gz | 20EPA011NSA-16_R2_001.fastq.gz |
| SRR32527843 | 20EPA011NSA_18 | ILLUMINA | NextSeq 550 | 20EPA011NSA-18_R1_001.fastq.gz | 20EPA011NSA-18_R2_001.fastq.gz |
| SRR32527842 | 20EPA011NSA_19 | ILLUMINA | NextSeq 550 | 20EPA011NSA-19_R1_001.fastq.gz | 20EPA011NSA-19_R2_001.fastq.gz |
| SRR32527841 | 20EPA011NSA_20 | ILLUMINA | NextSeq 550 | 20EPA011NSA-20_R1_001.fastq.gz | 20EPA011NSA-20_R2_001.fastq.gz |
| SRR32527840 | 20EPA012NSA_1 | ILLUMINA | NextSeq 550 | 20EPA012NSA-1_R1_001.fastq.gz | 20EPA012NSA-1_R2_001.fastq.gz |
| SRR32527838 | 20EPA012NSA_2 | ILLUMINA | NextSeq 550 | 20EPA012NSA-2_R1_001.fastq.gz | 20EPA012NSA-2_R2_001.fastq.gz |
| SRR32527837 | 20EPA012NSA_3 | ILLUMINA | NextSeq 550 | 20EPA012NSA-3_R1_001.fastq.gz | 20EPA012NSA-3_R2_001.fastq.gz |
| SRR32527836 | 20EPA012NSA_4 | ILLUMINA | NextSeq 550 | 20EPA012NSA-4_R1_001.fastq.gz | 20EPA012NSA-4_R2_001.fastq.gz |
| SRR32527835 | 20EPA012NSA_5 | ILLUMINA | NextSeq 550 | 20EPA012NSA-5_R1_001.fastq.gz | 20EPA012NSA-5_R2_001.fastq.gz |
| SRR32527834 | 20EPA012NSA_6 | ILLUMINA | NextSeq 550 | 20EPA012NSA-6_R1_001.fastq.gz | 20EPA012NSA-6_R2_001.fastq.gz |
| SRR32527833 | 20EPA012NSA_7 | ILLUMINA | NextSeq 550 | 20EPA012NSA-7_R1_001.fastq.gz | 20EPA012NSA-7_R2_001.fastq.gz |
| SRR32527832 | 20EPA012NSA_9 | ILLUMINA | NextSeq 550 | 20EPA012NSA-9_R1_001.fastq.gz | 20EPA012NSA-9_R2_001.fastq.gz |
| SRR32527831 | 20EPA012NSA_10 | ILLUMINA | NextSeq 550 | 20EPA012NSA-10_R1_001.fastq.gz | 20EPA012NSA-10_R2_001.fastq.gz |
| SRR32527830 | 20EPA012NSA_11 | ILLUMINA | NextSeq 550 | 20EPA012NSA-11_R1_001.fastq.gz | 20EPA012NSA-11_R2_001.fastq.gz |
| SRR32527829 | 20EPA012NSA_12 | ILLUMINA | NextSeq 550 | 20EPA012NSA-12_R1_001.fastq.gz | 20EPA012NSA-12_R2_001.fastq.gz |
| SRR32528019 | 20EPA012NSA_13 | ILLUMINA | NextSeq 550 | 20EPA012NSA-13_R1_001.fastq.gz | 20EPA012NSA-13_R2_001.fastq.gz |
| SRR32528018 | 20EPA012NSA_14 | ILLUMINA | NextSeq 550 | 20EPA012NSA-14_R1_001.fastq.gz | 20EPA012NSA-14_R2_001.fastq.gz |
| SRR32528017 | 20EPA012NSA_15 | ILLUMINA | NextSeq 550 | 20EPA012NSA-15_R1_001.fastq.gz | 20EPA012NSA-15_R2_001.fastq.gz |
| SRR32528016 | 20EPA012NSA_16 | ILLUMINA | NextSeq 550 | 20EPA012NSA-16_R1_001.fastq.gz | 20EPA012NSA-16_R2_001.fastq.gz |

| | | | | | |
|---|---|---|---|---|---|
| SRR32528015 | 20EPA012NSA_17 | ILLUMINA | NextSeq 550 | 20EPA012NSA-17_R1_001.fastq.gz | 20EPA012NSA-17_R2_001.fastq.gz |
| SRR32528014 | 20EPA012NSA_18 | ILLUMINA | NextSeq 550 | 20EPA012NSA-18_R1_001.fastq.gz | 20EPA012NSA-18_R2_001.fastq.gz |
| SRR32528013 | 20EPA012NSA_20 | ILLUMINA | NextSeq 550 | 20EPA012NSA-20_R1_001.fastq.gz | 20EPA012NSA-20_R2_001.fastq.gz |
| SRR32528012 | 22EPA044NSA_1 | ILLUMINA | NextSeq 550 | 22EPA044NSA-1_R1_001.fastq.gz | 22EPA044NSA-1_R2_001.fastq.gz |
| SRR32528011 | 22EPA044NSA_2 | ILLUMINA | NextSeq 550 | 22EPA044NSA-2_R1_001.fastq.gz | 22EPA044NSA-2_R2_001.fastq.gz |
| SRR32528010 | 22EPA044NSA_3 | ILLUMINA | NextSeq 550 | 22EPA044NSA-3_R1_001.fastq.gz | 22EPA044NSA-3_R2_001.fastq.gz |
| SRR32528008 | 22EPA044NSA_4 | ILLUMINA | NextSeq 550 | 22EPA044NSA-4_R1_001.fastq.gz | 22EPA044NSA-4_R2_001.fastq.gz |
| SRR32528007 | 22EPA044NSA_5 | ILLUMINA | NextSeq 550 | 22EPA044NSA-5_R1_001.fastq.gz | 22EPA044NSA-5_R2_001.fastq.gz |
| SRR32528006 | 22EPA044NSA_6 | ILLUMINA | NextSeq 550 | 22EPA044NSA-6_R1_001.fastq.gz | 22EPA044NSA-6_R2_001.fastq.gz |
| SRR32528005 | 22EPA044NSA_7 | ILLUMINA | NextSeq 550 | 22EPA044NSA-7_R1_001.fastq.gz | 22EPA044NSA-7_R2_001.fastq.gz |
| SRR32528004 | 22EPA044NSA_8 | ILLUMINA | NextSeq 550 | 22EPA044NSA-8_R1_001.fastq.gz | 22EPA044NSA-8_R2_001.fastq.gz |
| SRR32528003 | 22EPA044NSA_9 | ILLUMINA | NextSeq 550 | 22EPA044NSA-9_R1_001.fastq.gz | 22EPA044NSA-9_R2_001.fastq.gz |
| SRR32528002 | 22EPA044NSA_10 | ILLUMINA | NextSeq 550 | 22EPA044NSA-10_R1_001.fastq.gz | 22EPA044NSA-10_R2_001.fastq.gz |
| SRR32528001 | 22EPA044NSA_11 | ILLUMINA | NextSeq 550 | 22EPA044NSA-11_R1_001.fastq.gz | 22EPA044NSA-11_R2_001.fastq.gz |
| SRR32528000 | 22EPA044NSA_12 | ILLUMINA | NextSeq 550 | 22EPA044NSA-12_R1_001.fastq.gz | 22EPA044NSA-12_R2_001.fastq.gz |
| SRR32527999 | 22EPA044NSA_13 | ILLUMINA | NextSeq 550 | 22EPA044NSA-13_R1_001.fastq.gz | 22EPA044NSA-13_R2_001.fastq.gz |
| SRR32527997 | 22EPA044NSA_14 | ILLUMINA | NextSeq 550 | 22EPA044NSA-14_R1_001.fastq.gz | 22EPA044NSA-14_R2_001.fastq.gz |
| SRR32527996 | 22EPA044NSA_15 | ILLUMINA | NextSeq 550 | 22EPA044NSA-15_R1_001.fastq.gz | 22EPA044NSA-15_R2_001.fastq.gz |
| SRR32527995 | 22EPA044NSA_16 | ILLUMINA | NextSeq 550 | 22EPA044NSA-16_R1_001.fastq.gz | 22EPA044NSA-16_R2_001.fastq.gz |
| SRR32527994 | 22EPA044NSA_17 | ILLUMINA | NextSeq 550 | 22EPA044NSA-17_R1_001.fastq.gz | 22EPA044NSA-17_R2_001.fastq.gz |
| SRR32527993 | 22EPA044NSA_18 | ILLUMINA | NextSeq 550 | 22EPA044NSA-18_R1_001.fastq.gz | 22EPA044NSA-18_R2_001.fastq.gz |
| SRR32527992 | 22EPA044NSA_19 | ILLUMINA | NextSeq 550 | 22EPA044NSA-19_R1_001.fastq.gz | 22EPA044NSA-19_R2_001.fastq.gz |
| SRR32527991 | 22EPA044NSA_20 | ILLUMINA | NextSeq 550 | 22EPA044NSA-20_R1_001.fastq.gz | 22EPA044NSA-20_R2_001.fastq.gz |
| SRR32527990 | 22EPA051NSA_1 | ILLUMINA | NextSeq 550 | 22EPA051NSA-1_R1_001.fastq.gz | 22EPA051NSA-1_R2_001.fastq.gz |
| SRR32527989 | 22EPA051NSA_2 | ILLUMINA | NextSeq 550 | 22EPA051NSA-2_R1_001.fastq.gz | 22EPA051NSA-2_R2_001.fastq.gz |
| SRR32528043 | 22EPA051NSA_3 | ILLUMINA | NextSeq 550 | 22EPA051NSA-3_R1_001.fastq.gz | 22EPA051NSA-3_R2_001.fastq.gz |
| SRR32528041 | 22EPA051NSA_4 | ILLUMINA | NextSeq 550 | 22EPA051NSA-4_R1_001.fastq.gz | 22EPA051NSA-4_R2_001.fastq.gz |
| SRR32528040 | 22EPA051NSA_5 | ILLUMINA | NextSeq 550 | 22EPA051NSA-5_R1_001.fastq.gz | 22EPA051NSA-5_R2_001.fastq.gz |

| | | | | | |
|---|---|---|---|---|---|
| SRR32528039 | 22EPA051NSA_6 | ILLUMINA | NextSeq 550 | 22EPA051NSA-6_R1_001.fastq.gz | 22EPA051NSA-6_R2_001.fastq.gz |
| SRR32528038 | 22EPA051NSA_7 | ILLUMINA | NextSeq 550 | 22EPA051NSA-7_R1_001.fastq.gz | 22EPA051NSA-7_R2_001.fastq.gz |
| SRR32528037 | 22EPA051NSA_8 | ILLUMINA | NextSeq 550 | 22EPA051NSA-8_R1_001.fastq.gz | 22EPA051NSA-8_R2_001.fastq.gz |
| SRR32528036 | 22EPA051NSA_9 | ILLUMINA | NextSeq 550 | 22EPA051NSA-9_R1_001.fastq.gz | 22EPA051NSA-9_R2_001.fastq.gz |
| SRR32528035 | 22EPA051NSA_10 | ILLUMINA | NextSeq 550 | 22EPA051NSA-10_R1_001.fastq.gz | 22EPA051NSA-10_R2_001.fastq.gz |
| SRR32528034 | 22EPA051NSA_11 | ILLUMINA | NextSeq 550 | 22EPA051NSA-11_R1_001.fastq.gz | 22EPA051NSA-11_R2_001.fastq.gz |
| SRR32528033 | 22EPA051NSA_12 | ILLUMINA | NextSeq 550 | 22EPA051NSA-12_R1_001.fastq.gz | 22EPA051NSA-12_R2_001.fastq.gz |
| SRR32528032 | 22EPA051NSA_13 | ILLUMINA | NextSeq 550 | 22EPA051NSA-13_R1_001.fastq.gz | 22EPA051NSA-13_R2_001.fastq.gz |
| SRR32528030 | 22EPA051NSA_14 | ILLUMINA | NextSeq 550 | 22EPA051NSA-14_R1_001.fastq.gz | 22EPA051NSA-14_R2_001.fastq.gz |
| SRR32528029 | 22EPA051NSA_15 | ILLUMINA | NextSeq 550 | 22EPA051NSA-15_R1_001.fastq.gz | 22EPA051NSA-15_R2_001.fastq.gz |
| SRR32528028 | 22EPA051NSA_16 | ILLUMINA | NextSeq 550 | 22EPA051NSA-16_R1_001.fastq.gz | 22EPA051NSA-16_R2_001.fastq.gz |
| SRR32528027 | 22EPA051NSA_17 | ILLUMINA | NextSeq 550 | 22EPA051NSA-17_R1_001.fastq.gz | 22EPA051NSA-17_R2_001.fastq.gz |
| SRR32528026 | 22EPA051NSA_18 | ILLUMINA | NextSeq 550 | 22EPA051NSA-18_R1_001.fastq.gz | 22EPA051NSA-18_R2_001.fastq.gz |
| SRR32528025 | 22EPA051NSA_19 | ILLUMINA | NextSeq 550 | 22EPA051NSA-19_R1_001.fastq.gz | 22EPA051NSA-19_R2_001.fastq.gz |
| SRR32528024 | 22EPA051NSA_20 | ILLUMINA | NextSeq 550 | 22EPA051NSA-20_R1_001.fastq.gz | 22EPA051NSA-20_R2_001.fastq.gz |
| SRR32528023 | 22EPA053NSA_1 | ILLUMINA | NextSeq 550 | 22EPA053NSA-1_R1_001.fastq.gz | 22EPA053NSA-1_R2_001.fastq.gz |
| SRR32528022 | 22EPA053NSA_2 | ILLUMINA | NextSeq 550 | 22EPA053NSA-2_R1_001.fastq.gz | 22EPA053NSA-2_R2_001.fastq.gz |
| SRR32528021 | 22EPA053NSA_3 | ILLUMINA | NextSeq 550 | 22EPA053NSA-3_R1_001.fastq.gz | 22EPA053NSA-3_R2_001.fastq.gz |
| SRR32528136 | 22EPA053NSA_4 | ILLUMINA | NextSeq 550 | 22EPA053NSA-4_R1_001.fastq.gz | 22EPA053NSA-4_R2_001.fastq.gz |
| SRR32528135 | 22EPA053NSA_5 | ILLUMINA | NextSeq 550 | 22EPA053NSA-5_R1_001.fastq.gz | 22EPA053NSA-5_R2_001.fastq.gz |
| SRR32528134 | 22EPA053NSA_6 | ILLUMINA | NextSeq 550 | 22EPA053NSA-6_R1_001.fastq.gz | 22EPA053NSA-6_R2_001.fastq.gz |
| SRR32528133 | 22EPA053NSA_7 | ILLUMINA | NextSeq 550 | 22EPA053NSA-7_R1_001.fastq.gz | 22EPA053NSA-7_R2_001.fastq.gz |
| SRR32528132 | 22EPA053NSA_8 | ILLUMINA | NextSeq 550 | 22EPA053NSA-8_R1_001.fastq.gz | 22EPA053NSA-8_R2_001.fastq.gz |
| SRR32528131 | 22EPA053NSA_9 | ILLUMINA | NextSeq 550 | 22EPA053NSA-9_R1_001.fastq.gz | 22EPA053NSA-9_R2_001.fastq.gz |
| SRR32528130 | 22EPA053NSA_10 | ILLUMINA | NextSeq 550 | 22EPA053NSA-10_R1_001.fastq.gz | 22EPA053NSA-10_R2_001.fastq.gz |
| SRR32528129 | 22EPA053NSA_11 | ILLUMINA | NextSeq 550 | 22EPA053NSA-11_R1_001.fastq.gz | 22EPA053NSA-11_R2_001.fastq.gz |
| SRR32528128 | 22EPA053NSA_12 | ILLUMINA | NextSeq 550 | 22EPA053NSA-12_R1_001.fastq.gz | 22EPA053NSA-12_R2_001.fastq.gz |
| SRR32528127 | 22EPA053NSA_13 | ILLUMINA | NextSeq 550 | 22EPA053NSA-13_R1_001.fastq.gz | 22EPA053NSA-13_R2_001.fastq.gz |

| | | | | | |
|---|---|---|---|---|---|
| SRR32528125 | 22EPA053NSA_14 | ILLUMINA | NextSeq 550 | 22EPA053NSA-14_R1_001.fastq.gz | 22EPA053NSA-14_R2_001.fastq.gz |
| SRR32528124 | 22EPA053NSA_15 | ILLUMINA | NextSeq 550 | 22EPA053NSA-15_R1_001.fastq.gz | 22EPA053NSA-15_R2_001.fastq.gz |
| SRR32528123 | 22EPA053NSA_16 | ILLUMINA | NextSeq 550 | 22EPA053NSA-16_R1_001.fastq.gz | 22EPA053NSA-16_R2_001.fastq.gz |
| SRR32528122 | 22EPA053NSA_17 | ILLUMINA | NextSeq 550 | 22EPA053NSA-17_R1_001.fastq.gz | 22EPA053NSA-17_R2_001.fastq.gz |
| SRR32528121 | 22EPA053NSA_18 | ILLUMINA | NextSeq 550 | 22EPA053NSA-18_R1_001.fastq.gz | 22EPA053NSA-18_R2_001.fastq.gz |
| SRR32528120 | 22EPA053NSA_19 | ILLUMINA | NextSeq 550 | 22EPA053NSA-19_R1_001.fastq.gz | 22EPA053NSA-19_R2_001.fastq.gz |
| SRR32528119 | 22EPA053NSA_20 | ILLUMINA | NextSeq 550 | 22EPA053NSA-20_R1_001.fastq.gz | 22EPA053NSA-20_R2_001.fastq.gz |
| SRR32528118 | 22EPA055NSA_1 | ILLUMINA | NextSeq 550 | 22EPA055NSA-1_R1_001.fastq.gz | 22EPA055NSA-1_R2_001.fastq.gz |
| SRR32528117 | 22EPA055NSA_2 | ILLUMINA | NextSeq 550 | 22EPA055NSA-2_R1_001.fastq.gz | 22EPA055NSA-2_R2_001.fastq.gz |
| SRR32528116 | 22EPA055NSA_3 | ILLUMINA | NextSeq 550 | 22EPA055NSA-3_R1_001.fastq.gz | 22EPA055NSA-3_R2_001.fastq.gz |
| SRR32528114 | 22EPA055NSA_4 | ILLUMINA | NextSeq 550 | 22EPA055NSA-4_R1_001.fastq.gz | 22EPA055NSA-4_R2_001.fastq.gz |
| SRR32528113 | 22EPA055NSA_5 | ILLUMINA | NextSeq 550 | 22EPA055NSA-5_R1_001.fastq.gz | 22EPA055NSA-5_R2_001.fastq.gz |
| SRR32528112 | 22EPA055NSA_6 | ILLUMINA | NextSeq 550 | 22EPA055NSA-6_R1_001.fastq.gz | 22EPA055NSA-6_R2_001.fastq.gz |
| SRR32528111 | 22EPA055NSA_7 | ILLUMINA | NextSeq 550 | 22EPA055NSA-7_R1_001.fastq.gz | 22EPA055NSA-7_R2_001.fastq.gz |
| SRR32528110 | 22EPA055NSA_8 | ILLUMINA | NextSeq 550 | 22EPA055NSA-8_R1_001.fastq.gz | 22EPA055NSA-8_R2_001.fastq.gz |
| SRR32528109 | 22EPA055NSA_9 | ILLUMINA | NextSeq 550 | 22EPA055NSA-9_R1_001.fastq.gz | 22EPA055NSA-9_R2_001.fastq.gz |
| SRR32528108 | 22EPA055NSA_10 | ILLUMINA | NextSeq 550 | 22EPA055NSA-10_R1_001.fastq.gz | 22EPA055NSA-10_R2_001.fastq.gz |
| SRR32527988 | 22EPA055NSA_11 | ILLUMINA | NextSeq 550 | 22EPA055NSA-11_R1_001.fastq.gz | 22EPA055NSA-11_R2_001.fastq.gz |
| SRR32527987 | 22EPA055NSA_12 | ILLUMINA | NextSeq 550 | 22EPA055NSA-12_R1_001.fastq.gz | 22EPA055NSA-12_R2_001.fastq.gz |
| SRR32527986 | 22EPA055NSA_13 | ILLUMINA | NextSeq 550 | 22EPA055NSA-13_R1_001.fastq.gz | 22EPA055NSA-13_R2_001.fastq.gz |
| SRR32527984 | 22EPA055NSA_14 | ILLUMINA | NextSeq 550 | 22EPA055NSA-14_R1_001.fastq.gz | 22EPA055NSA-14_R2_001.fastq.gz |
| SRR32527983 | 22EPA055NSA_15 | ILLUMINA | NextSeq 550 | 22EPA055NSA-15_R1_001.fastq.gz | 22EPA055NSA-15_R2_001.fastq.gz |
| SRR32527982 | 22EPA055NSA_16 | ILLUMINA | NextSeq 550 | 22EPA055NSA-16_R1_001.fastq.gz | 22EPA055NSA-16_R2_001.fastq.gz |
| SRR32527981 | 22EPA055NSA_17 | ILLUMINA | NextSeq 550 | 22EPA055NSA-17_R1_001.fastq.gz | 22EPA055NSA-17_R2_001.fastq.gz |
| SRR32527980 | 22EPA055NSA_18 | ILLUMINA | NextSeq 550 | 22EPA055NSA-18_R1_001.fastq.gz | 22EPA055NSA-18_R2_001.fastq.gz |
| SRR32527979 | 22EPA055NSA_19 | ILLUMINA | NextSeq 550 | 22EPA055NSA-19_R1_001.fastq.gz | 22EPA055NSA-19_R2_001.fastq.gz |
| SRR32527978 | 22EPA055NSA_20 | ILLUMINA | NextSeq 550 | 22EPA055NSA-20_R1_001.fastq.gz | 22EPA055NSA-20_R2_001.fastq.gz |
| SRR32527977 | 22EPA058NSA_1 | ILLUMINA | NextSeq 550 | 22EPA058NSA-1_R1_001.fastq.gz | 22EPA058NSA-1_R2_001.fastq.gz |

| | | | | | |
|---|---|---|---|---|---|
| SRR32527976 | 22EPA058NSA_2 | ILLUMINA | NextSeq 550 | 22EPA058NSA-2_R1_001.fastq.gz | 22EPA058NSA-2_R2_001.fastq.gz |
| SRR32527975 | 22EPA058NSA_3 | ILLUMINA | NextSeq 550 | 22EPA058NSA-3_R1_001.fastq.gz | 22EPA058NSA-3_R2_001.fastq.gz |
| SRR32527973 | 22EPA058NSA_4 | ILLUMINA | NextSeq 550 | 22EPA058NSA-4_R1_001.fastq.gz | 22EPA058NSA-4_R2_001.fastq.gz |
| SRR32527972 | 22EPA058NSA_5 | ILLUMINA | NextSeq 550 | 22EPA058NSA-5_R1_001.fastq.gz | 22EPA058NSA-5_R2_001.fastq.gz |
| SRR32527971 | 22EPA058NSA_6 | ILLUMINA | NextSeq 550 | 22EPA058NSA-6_R1_001.fastq.gz | 22EPA058NSA-6_R2_001.fastq.gz |
| SRR32527970 | 22EPA058NSA_7 | ILLUMINA | NextSeq 550 | 22EPA058NSA-7_R1_001.fastq.gz | 22EPA058NSA-7_R2_001.fastq.gz |
| SRR32527969 | 22EPA058NSA_8 | ILLUMINA | NextSeq 550 | 22EPA058NSA-8_R1_001.fastq.gz | 22EPA058NSA-8_R2_001.fastq.gz |
| SRR32527968 | 22EPA058NSA_9 | ILLUMINA | NextSeq 550 | 22EPA058NSA-9_R1_001.fastq.gz | 22EPA058NSA-9_R2_001.fastq.gz |
| SRR32527967 | 22EPA058NSA_10 | ILLUMINA | NextSeq 550 | 22EPA058NSA-10_R1_001.fastq.gz | 22EPA058NSA-10_R2_001.fastq.gz |
| SRR32527966 | 22EPA058NSA_11 | ILLUMINA | NextSeq 550 | 22EPA058NSA-11_R1_001.fastq.gz | 22EPA058NSA-11_R2_001.fastq.gz |
| SRR32527965 | 22EPA058NSA_12 | ILLUMINA | NextSeq 550 | 22EPA058NSA-12_R1_001.fastq.gz | 22EPA058NSA-12_R2_001.fastq.gz |
| SRR32527964 | 22EPA058NSA_13 | ILLUMINA | NextSeq 550 | 22EPA058NSA-13_R1_001.fastq.gz | 22EPA058NSA-13_R2_001.fastq.gz |
| SRR32527962 | 22EPA058NSA_14 | ILLUMINA | NextSeq 550 | 22EPA058NSA-14_R1_001.fastq.gz | 22EPA058NSA-14_R2_001.fastq.gz |
| SRR32527961 | 22EPA058NSA_15 | ILLUMINA | NextSeq 550 | 22EPA058NSA-15_R1_001.fastq.gz | 22EPA058NSA-15_R2_001.fastq.gz |
| SRR32527960 | 22EPA058NSA_16 | ILLUMINA | NextSeq 550 | 22EPA058NSA-16_R1_001.fastq.gz | 22EPA058NSA-16_R2_001.fastq.gz |
| SRR32527959 | 22EPA058NSA_17 | ILLUMINA | NextSeq 550 | 22EPA058NSA-17_R1_001.fastq.gz | 22EPA058NSA-17_R2_001.fastq.gz |
| SRR32527958 | 22EPA058NSA_18 | ILLUMINA | NextSeq 550 | 22EPA058NSA-18_R1_001.fastq.gz | 22EPA058NSA-18_R2_001.fastq.gz |
| SRR32527957 | 22EPA058NSA_19 | ILLUMINA | NextSeq 550 | 22EPA058NSA-19_R1_001.fastq.gz | 22EPA058NSA-19_R2_001.fastq.gz |
| SRR32527924 | 22EPA058NSA_20 | ILLUMINA | NextSeq 550 | 22EPA058NSA-20_R1_001.fastq.gz | 22EPA058NSA-20_R2_001.fastq.gz |

Table S3. Accession numbers for the Oxford Nanopore reads used in study

| Accession | Sample_name | Platform | Instrument_model | Filename |
|---|---|---|---|---|
| SRR32527923 | 20EPA002NSA_1 | ONT | MinION | 20EPA002NSA_1.fastq |
| SRR32527922 | 20EPA002NSA_2 | ONT | MinION | 20EPA002NSA_2.fastq |
| SRR32527921 | 20EPA002NSA_3 | ONT | MinION | 20EPA002NSA_3.fastq |
| SRR32527919 | 20EPA002NSA_4 | ONT | MinION | 20EPA002NSA_4.fastq |
| SRR32527918 | 20EPA002NSA_5 | ONT | MinION | 20EPA002NSA_5.fastq |
| SRR32527917 | 20EPA002NSA_6 | ONT | MinION | 20EPA002NSA_6.fastq |
| SRR32527916 | 20EPA002NSA_7 | ONT | MinION | 20EPA002NSA_7.fastq |
| SRR32527915 | 20EPA002NSA_8 | ONT | MinION | 20EPA002NSA_8.fastq |
| SRR32527914 | 20EPA002NSA_9 | ONT | MinION | 20EPA002NSA_9.fastq |
| SRR32527913 | 20EPA002NSA_10 | ONT | MinION | 20EPA002NSA_10.fastq |
| SRR32527912 | 20EPA002NSA_12 | ONT | MinION | 20EPA002NSA_12.fastq |
| SRR32527911 | 20EPA002NSA_13 | ONT | MinION | 20EPA002NSA_13.fastq |
| SRR32527910 | 20EPA002NSA_14 | ONT | MinION | 20EPA002NSA_14.fastq |
| SRR32527908 | 20EPA002NSA_15 | ONT | MinION | 20EPA002NSA_15.fastq |
| SRR32527907 | 20EPA002NSA_16 | ONT | MinION | 20EPA002NSA_16.fastq |
| SRR32527906 | 20EPA002NSA_17 | ONT | MinION | 20EPA002NSA_17.fastq |
| SRR32527905 | 20EPA002NSA_18 | ONT | MinION | 20EPA002NSA_18.fastq |
| SRR32527904 | 20EPA002NSA_19 | ONT | MinION | 20EPA002NSA_19.fastq |
| SRR32527903 | 20EPA002NSA_20 | ONT | MinION | 20EPA002NSA_20.fastq |
| SRR32527902 | 20EPA011NSA_1 | ONT | MinION | 20EPA011NSA_1.fastq |
| SRR32527901 | 20EPA011NSA_2 | ONT | MinION | 20EPA011NSA_2.fastq |
| SRR32527900 | 20EPA011NSA_3 | ONT | MinION | 20EPA011NSA_3.fastq |
| SRR32527899 | 20EPA011NSA_4 | ONT | MinION | 20EPA011NSA_4.fastq |
| SRR32527897 | 20EPA011NSA_5 | ONT | MinION | 20EPA011NSA_5.fastq |
| SRR32527896 | 20EPA011NSA_6 | ONT | MinION | 20EPA011NSA_6.fastq |
| SRR32527895 | 20EPA011NSA_7 | ONT | MinION | 20EPA011NSA_7.fastq |
| SRR32527894 | 20EPA011NSA_8 | ONT | MinION | 20EPA011NSA_8.fastq |
| SRR32527893 | 20EPA011NSA_9 | ONT | MinION | 20EPA011NSA_9.fastq |
| SRR32528107 | 20EPA011NSA_10 | ONT | MinION | 20EPA011NSA_10.fastq |
| SRR32528106 | 20EPA011NSA_11 | ONT | MinION | 20EPA011NSA_11.fastq |
| SRR32528105 | 20EPA011NSA_12 | ONT | MinION | 20EPA011NSA_12.fastq |
| SRR32528104 | 20EPA011NSA_13 | ONT | MinION | 20EPA011NSA_13.fastq |
| SRR32528103 | 20EPA011NSA_14 | ONT | MinION | 20EPA011NSA_14.fastq |
| SRR32528101 | 20EPA011NSA_15 | ONT | MinION | 20EPA011NSA_15.fastq |
| SRR32528100 | 20EPA011NSA_16 | ONT | MinION | 20EPA011NSA_16.fastq |
| SRR32528099 | 20EPA011NSA_18 | ONT | MinION | 20EPA011NSA_18.fastq |
| SRR32528098 | 20EPA011NSA_19 | ONT | MinION | 20EPA011NSA_19.fastq |
| SRR32528097 | 20EPA011NSA_20 | ONT | MinION | 20EPA011NSA_20.fastq |
| SRR32528096 | 20EPA012NSA_1 | ONT | MinION | 20EPA012NSA_1.fastq |
| SRR32528095 | 20EPA012NSA_2 | ONT | MinION | 20EPA012NSA_2.fastq |
| SRR32528094 | 20EPA012NSA_3 | ONT | MinION | 20EPA012NSA_3.fastq |
| SRR32528093 | 20EPA012NSA_4 | ONT | MinION | 20EPA012NSA_4.fastq |

| | | | | |
|---|---|---|---|---|
| SRR32528092 | 20EPA012NSA_5 | ONT | MinION | 20EPA012NSA_5.fastq |
| SRR32528089 | 20EPA012NSA_6 | ONT | MinION | 20EPA012NSA_6.fastq |
| SRR32528088 | 20EPA012NSA_7 | ONT | MinION | 20EPA012NSA_7.fastq |
| SRR32528087 | 20EPA012NSA_9 | ONT | MinION | 20EPA012NSA_9.fastq |
| SRR32528086 | 20EPA012NSA_10 | ONT | MinION | 20EPA012NSA_10.fastq |
| SRR32528085 | 20EPA012NSA_11 | ONT | MinION | 20EPA012NSA_11.fastq |
| SRR32528084 | 20EPA012NSA_12 | ONT | MinION | 20EPA012NSA_12.fastq |
| SRR32528083 | 20EPA012NSA_13 | ONT | MinION | 20EPA012NSA_13.fastq |
| SRR32528082 | 20EPA012NSA_14 | ONT | MinION | 20EPA012NSA_14.fastq |
| SRR32528081 | 20EPA012NSA_15 | ONT | MinION | 20EPA012NSA_15.fastq |
| SRR32528080 | 20EPA012NSA_16 | ONT | MinION | 20EPA012NSA_16.fastq |
| SRR32528078 | 20EPA012NSA_17 | ONT | MinION | 20EPA012NSA_17.fastq |
| SRR32528077 | 20EPA012NSA_18 | ONT | MinION | 20EPA012NSA_18.fastq |
| SRR32528076 | 20EPA012NSA_20 | ONT | MinION | 20EPA012NSA_20.fastq |
| SRR32527956 | 22EPA044NSA_1 | ONT | MinION | 22EPA044NSA_1.fastq |
| SRR32527955 | 22EPA044NSA_2 | ONT | MinION | 22EPA044NSA_2.fastq |
| SRR32527954 | 22EPA044NSA_3 | ONT | MinION | 22EPA044NSA_3.fastq |
| SRR32527953 | 22EPA044NSA_4 | ONT | MinION | 22EPA044NSA_4.fastq |
| SRR32527952 | 22EPA044NSA_5 | ONT | MinION | 22EPA044NSA_5.fastq |
| SRR32527951 | 22EPA044NSA_6 | ONT | MinION | 22EPA044NSA_6.fastq |
| SRR32527950 | 22EPA044NSA_7 | ONT | MinION | 22EPA044NSA_7.fastq |
| SRR32527948 | 22EPA044NSA_8 | ONT | MinION | 22EPA044NSA_8.fastq |
| SRR32527947 | 22EPA044NSA_9 | ONT | MinION | 22EPA044NSA_9.fastq |
| SRR32527946 | 22EPA044NSA_10 | ONT | MinION | 22EPA044NSA_10.fastq |
| SRR32527945 | 22EPA044NSA_11 | ONT | MinION | 22EPA044NSA_11.fastq |
| SRR32527944 | 22EPA044NSA_12 | ONT | MinION | 22EPA044NSA_12.fastq |
| SRR32527943 | 22EPA044NSA_13 | ONT | MinION | 22EPA044NSA_13.fastq |
| SRR32527942 | 22EPA044NSA_14 | ONT | MinION | 22EPA044NSA_14.fastq |
| SRR32527941 | 22EPA044NSA_15 | ONT | MinION | 22EPA044NSA_15.fastq |
| SRR32527940 | 22EPA044NSA_16 | ONT | MinION | 22EPA044NSA_16.fastq |
| SRR32527939 | 22EPA044NSA_17 | ONT | MinION | 22EPA044NSA_17.fastq |
| SRR32527937 | 22EPA044NSA_18 | ONT | MinION | 22EPA044NSA_18.fastq |
| SRR32527936 | 22EPA044NSA_19 | ONT | MinION | 22EPA044NSA_19.fastq |
| SRR32527935 | 22EPA044NSA_20 | ONT | MinION | 22EPA044NSA_20.fastq |
| SRR32527934 | 22EPA051NSA_1 | ONT | MinION | 22EPA051NSA_1.fastq |
| SRR32527933 | 22EPA051NSA_2 | ONT | MinION | 22EPA051NSA_2.fastq |
| SRR32527932 | 22EPA051NSA_3 | ONT | MinION | 22EPA051NSA_3.fastq |
| SRR32527931 | 22EPA051NSA_4 | ONT | MinION | 22EPA051NSA_4.fastq |
| SRR32527930 | 22EPA051NSA_5 | ONT | MinION | 22EPA051NSA_5.fastq |
| SRR32527929 | 22EPA051NSA_6 | ONT | MinION | 22EPA051NSA_6.fastq |
| SRR32527928 | 22EPA051NSA_7 | ONT | MinION | 22EPA051NSA_7.fastq |
| SRR32527926 | 22EPA051NSA_8 | ONT | MinION | 22EPA051NSA_8.fastq |
| SRR32527925 | 22EPA051NSA_9 | ONT | MinION | 22EPA051NSA_9.fastq |
| SRR32527892 | 22EPA051NSA_10 | ONT | MinION | 22EPA051NSA_10.fastq |

| | | | | |
|---|---|---|---|---|
| SRR32527891 | 22EPA051NSA_11 | ONT | MinION | 22EPA051NSA_11.fastq |
| SRR32527890 | 22EPA051NSA_12 | ONT | MinION | 22EPA051NSA_12.fastq |
| SRR32527889 | 22EPA051NSA_13 | ONT | MinION | 22EPA051NSA_13.fastq |
| SRR32527888 | 22EPA051NSA_14 | ONT | MinION | 22EPA051NSA_14.fastq |
| SRR32527887 | 22EPA051NSA_15 | ONT | MinION | 22EPA051NSA_15.fastq |
| SRR32527886 | 22EPA051NSA_16 | ONT | MinION | 22EPA051NSA_16.fastq |
| SRR32527885 | 22EPA051NSA_17 | ONT | MinION | 22EPA051NSA_17.fastq |
| SRR32527883 | 22EPA051NSA_18 | ONT | MinION | 22EPA051NSA_18.fastq |
| SRR32527882 | 22EPA051NSA_19 | ONT | MinION | 22EPA051NSA_19.fastq |
| SRR32527881 | 22EPA051NSA_20 | ONT | MinION | 22EPA051NSA_20.fastq |
| SRR32527880 | 22EPA053NSA_1 | ONT | MinION | 22EPA053NSA_1.fastq |
| SRR32527879 | 22EPA053NSA_2 | ONT | MinION | 22EPA053NSA_2.fastq |
| SRR32527878 | 22EPA053NSA_3 | ONT | MinION | 22EPA053NSA_3.fastq |
| SRR32527877 | 22EPA053NSA_4 | ONT | MinION | 22EPA053NSA_4.fastq |
| SRR32527876 | 22EPA053NSA_5 | ONT | MinION | 22EPA053NSA_5.fastq |
| SRR32527875 | 22EPA053NSA_6 | ONT | MinION | 22EPA053NSA_6.fastq |
| SRR32527874 | 22EPA053NSA_7 | ONT | MinION | 22EPA053NSA_7.fastq |
| SRR32527872 | 22EPA053NSA_8 | ONT | MinION | 22EPA053NSA_8.fastq |
| SRR32527871 | 22EPA053NSA_9 | ONT | MinION | 22EPA053NSA_9.fastq |
| SRR32527870 | 22EPA053NSA_10 | ONT | MinION | 22EPA053NSA_10.fastq |
| SRR32527869 | 22EPA053NSA_11 | ONT | MinION | 22EPA053NSA_11.fastq |
| SRR32527868 | 22EPA053NSA_12 | ONT | MinION | 22EPA053NSA_12.fastq |
| SRR32527867 | 22EPA053NSA_13 | ONT | MinION | 22EPA053NSA_13.fastq |
| SRR32527866 | 22EPA053NSA_14 | ONT | MinION | 22EPA053NSA_14.fastq |
| SRR32527865 | 22EPA053NSA_15 | ONT | MinION | 22EPA053NSA_15.fastq |
| SRR32527864 | 22EPA053NSA_16 | ONT | MinION | 22EPA053NSA_16.fastq |
| SRR32527863 | 22EPA053NSA_17 | ONT | MinION | 22EPA053NSA_17.fastq |
| SRR32527861 | 22EPA053NSA_18 | ONT | MinION | 22EPA053NSA_18.fastq |
| SRR32528075 | 22EPA053NSA_19 | ONT | MinION | 22EPA053NSA_19.fastq |
| SRR32528074 | 22EPA053NSA_20 | ONT | MinION | 22EPA053NSA_20.fastq |
| SRR32528073 | 22EPA055NSA_1 | ONT | MinION | 22EPA055NSA_1.fastq |
| SRR32528072 | 22EPA055NSA_2 | ONT | MinION | 22EPA055NSA_2.fastq |
| SRR32528071 | 22EPA055NSA_3 | ONT | MinION | 22EPA055NSA_3.fastq |
| SRR32528070 | 22EPA055NSA_4 | ONT | MinION | 22EPA055NSA_4.fastq |
| SRR32528069 | 22EPA055NSA_5 | ONT | MinION | 22EPA055NSA_5.fastq |
| SRR32528068 | 22EPA055NSA_6 | ONT | MinION | 22EPA055NSA_6.fastq |
| SRR32528067 | 22EPA055NSA_7 | ONT | MinION | 22EPA055NSA_7.fastq |
| SRR32528065 | 22EPA055NSA_8 | ONT | MinION | 22EPA055NSA_8.fastq |
| SRR32528064 | 22EPA055NSA_9 | ONT | MinION | 22EPA055NSA_9.fastq |
| SRR32528063 | 22EPA055NSA_10 | ONT | MinION | 22EPA055NSA_10.fastq |
| SRR32528062 | 22EPA055NSA_11 | ONT | MinION | 22EPA055NSA_11.fastq |
| SRR32528061 | 22EPA055NSA_12 | ONT | MinION | 22EPA055NSA_12.fastq |
| SRR32528060 | 22EPA055NSA_13 | ONT | MinION | 22EPA055NSA_13.fastq |
| SRR32528059 | 22EPA055NSA_14 | ONT | MinION | 22EPA055NSA_14.fastq |

| | | | | |
|---|---|---|---|---|
| SRR32528058 | 22EPA055NSA_15 | ONT | MinION | 22EPA055NSA_15.fastq |
| SRR32528057 | 22EPA055NSA_16 | ONT | MinION | 22EPA055NSA_16.fastq |
| SRR32528056 | 22EPA055NSA_17 | ONT | MinION | 22EPA055NSA_17.fastq |
| SRR32528054 | 22EPA055NSA_18 | ONT | MinION | 22EPA055NSA_18.fastq |
| SRR32528053 | 22EPA055NSA_19 | ONT | MinION | 22EPA055NSA_19.fastq |
| SRR32528052 | 22EPA055NSA_20 | ONT | MinION | 22EPA055NSA_20.fastq |
| SRR32528051 | 22EPA058NSA_1 | ONT | MinION | 22EPA058NSA_1.fastq |
| SRR32528050 | 22EPA058NSA_2 | ONT | MinION | 22EPA058NSA_2.fastq |
| SRR32528049 | 22EPA058NSA_3 | ONT | MinION | 22EPA058NSA_3.fastq |
| SRR32528048 | 22EPA058NSA_4 | ONT | MinION | 22EPA058NSA_4.fastq |
| SRR32528047 | 22EPA058NSA_5 | ONT | MinION | 22EPA058NSA_5.fastq |
| SRR32528046 | 22EPA058NSA_6 | ONT | MinION | 22EPA058NSA_6.fastq |
| SRR32528045 | 22EPA058NSA_7 | ONT | MinION | 22EPA058NSA_7.fastq |
| SRR32527859 | 22EPA058NSA_8 | ONT | MinION | 22EPA058NSA_8.fastq |
| SRR32527858 | 22EPA058NSA_9 | ONT | MinION | 22EPA058NSA_9.fastq |
| SRR32527857 | 22EPA058NSA_10 | ONT | MinION | 22EPA058NSA_10.fastq |
| SRR32527856 | 22EPA058NSA_11 | ONT | MinION | 22EPA058NSA_11.fastq |
| SRR32527855 | 22EPA058NSA_14 | ONT | MinION | 22EPA058NSA_14.fastq |
| SRR32527854 | 22EPA058NSA_15 | ONT | MinION | 22EPA058NSA_15.fastq |
| SRR32527853 | 22EPA058NSA_16 | ONT | MinION | 22EPA058NSA_16.fastq |
| SRR32527852 | 22EPA058NSA_17 | ONT | MinION | 22EPA058NSA_17.fastq |
| SRR32527851 | 22EPA058NSA_18 | ONT | MinION | 22EPA058NSA_18.fastq |
| SRR32527850 | 22EPA058NSA_19 | ONT | MinION | 22EPA058NSA_19.fastq |
| SRR32527848 | 22EPA058NSA_20 | ONT | MinION | 22EPA058NSA_20.fastq |

Table S4. Accession numbers for the hybrid assemblies used in the study

| Accession | Sample_name | Serovar |
|---|---|---|
| SAMN47165107 | 20EPA002NSA_1 | Java |
| SAMN47165108 | 20EPA002NSA_2 | Java |
| SAMN47165109 | 20EPA002NSA_3 | Java |
| SAMN47165110 | 20EPA002NSA_4 | Java |
| SAMN47165111 | 20EPA002NSA_5 | Java |
| SAMN47165112 | 20EPA002NSA_6 | Java |
| SAMN47165113 | 20EPA002NSA_7 | Java |
| SAMN47165114 | 20EPA002NSA_8 | Java |
| SAMN47165115 | 20EPA002NSA_9 | Java |
| SAMN47165116 | 20EPA002NSA_10 | Java |
| SAMN47165117 | 20EPA002NSA_11 | Java |
| SAMN47165118 | 20EPA002NSA_12 | Java |
| SAMN47165119 | 20EPA002NSA_13 | Java |
| SAMN47165120 | 20EPA002NSA_14 | Java |
| SAMN47165121 | 20EPA002NSA_15 | Java |
| SAMN47165122 | 20EPA002NSA_16 | Java |
| SAMN47165123 | 20EPA002NSA_17 | Java |
| SAMN47165124 | 20EPA002NSA_18 | Java |
| SAMN47165125 | 20EPA002NSA_19 | Java |
| SAMN47165126 | 20EPA002NSA_20 | Java |
| SAMN47165127 | 20EPA011NSA_1 | Java |
| SAMN47165128 | 20EPA011NSA_2 | Java |
| SAMN47165129 | 20EPA011NSA_3 | Java |
| SAMN47165130 | 20EPA011NSA_4 | Java |
| SAMN47165131 | 20EPA011NSA_5 | Java |
| SAMN47165132 | 20EPA011NSA_6 | Java |
| SAMN47165133 | 20EPA011NSA_7 | Java |
| SAMN47165134 | 20EPA011NSA_8 | Java |
| SAMN47165135 | 20EPA011NSA_9 | Java |
| SAMN47165136 | 20EPA011NSA_10 | Java |
| SAMN47165137 | 20EPA011NSA_11 | Java |
| SAMN47165138 | 20EPA011NSA_12 | Java |
| SAMN47165139 | 20EPA011NSA_13 | Java |
| SAMN47165140 | 20EPA011NSA_14 | Java |
| SAMN47165141 | 20EPA011NSA_15 | Java |
| SAMN47165142 | 20EPA011NSA_16 | Java |
| SAMN47165143 | 20EPA011NSA_18 | Java |
| SAMN47165144 | 20EPA011NSA_19 | Java |
| SAMN47165145 | 20EPA011NSA_20 | Java |
| SAMN47165146 | 20EPA012NSA_1 | Infantis |
| SAMN47165147 | 20EPA012NSA_2 | Infantis |
| SAMN47165148 | 20EPA012NSA_3 | Infantis |

| | | |
|---|---|---|
| SAMN47165149 | 20EPA012NSA_4 | Infantis |
| SAMN47165150 | 20EPA012NSA_5 | Infantis |
| SAMN47165151 | 20EPA012NSA_6 | Infantis |
| SAMN47165152 | 20EPA012NSA_7 | Infantis |
| SAMN47165153 | 20EPA012NSA_9 | Infantis |
| SAMN47165154 | 20EPA012NSA_10 | Infantis |
| SAMN47165155 | 20EPA012NSA_11 | Infantis |
| SAMN47165156 | 20EPA012NSA_12 | Infantis |
| SAMN47165157 | 20EPA012NSA_13 | Infantis |
| SAMN47165158 | 20EPA012NSA_14 | Infantis |
| SAMN47165159 | 20EPA012NSA_15 | Infantis |
| SAMN47165160 | 20EPA012NSA_16 | Infantis |
| SAMN47165161 | 20EPA012NSA_17 | Infantis |
| SAMN47165162 | 20EPA012NSA_18 | Infantis |
| SAMN47165163 | 20EPA012NSA_20 | Infantis |
| SAMN47165164 | 22EPA044NSA_1 | Typhimurium |
| SAMN47165165 | 22EPA044NSA_2 | Typhimurium |
| SAMN47165166 | 22EPA044NSA_3 | Typhimurium |
| SAMN47165167 | 22EPA044NSA_4 | Typhimurium |
| SAMN47165168 | 22EPA044NSA_5 | Typhimurium |
| SAMN47165169 | 22EPA044NSA_6 | Typhimurium |
| SAMN47165170 | 22EPA044NSA_7 | Typhimurium |
| SAMN47165171 | 22EPA044NSA_8 | Typhimurium |
| SAMN47165172 | 22EPA044NSA_9 | Typhimurium |
| SAMN47165173 | 22EPA044NSA_10 | Typhimurium |
| SAMN47165174 | 22EPA044NSA_11 | Typhimurium |
| SAMN47165175 | 22EPA044NSA_12 | Typhimurium |
| SAMN47165176 | 22EPA044NSA_13 | Typhimurium |
| SAMN47165177 | 22EPA044NSA_14 | Typhimurium |
| SAMN47165178 | 22EPA044NSA_15 | Typhimurium |
| SAMN47165179 | 22EPA044NSA_16 | Typhimurium |
| SAMN47165180 | 22EPA044NSA_17 | Typhimurium |
| SAMN47165181 | 22EPA044NSA_18 | Typhimurium |
| SAMN47165182 | 22EPA044NSA_19 | Typhimurium |
| SAMN47165183 | 22EPA044NSA_20 | Typhimurium |
| SAMN47165184 | 22EPA051NSA_1 | Enteritidis |
| SAMN47165185 | 22EPA051NSA_2 | Enteritidis |
| SAMN47165186 | 22EPA051NSA_3 | Enteritidis |
| SAMN47165187 | 22EPA051NSA_4 | Enteritidis |
| SAMN47165188 | 22EPA051NSA_5 | Enteritidis |
| SAMN47165189 | 22EPA051NSA_6 | Enteritidis |
| SAMN47165190 | 22EPA051NSA_7 | Enteritidis |
| SAMN47165191 | 22EPA051NSA_8 | Enteritidis |
| SAMN47165192 | 22EPA051NSA_9 | Enteritidis |

| | | |
|---|---|---|
| SAMN47165193 | 22EPA051NSA_10 | Enteritidis |
| SAMN47165194 | 22EPA051NSA_11 | Enteritidis |
| SAMN47165195 | 22EPA051NSA_12 | Enteritidis |
| SAMN47165196 | 22EPA051NSA_13 | Enteritidis |
| SAMN47165197 | 22EPA051NSA_14 | Enteritidis |
| SAMN47165198 | 22EPA051NSA_15 | Enteritidis |
| SAMN47165199 | 22EPA051NSA_16 | Enteritidis |
| SAMN47165200 | 22EPA051NSA_17 | Enteritidis |
| SAMN47165201 | 22EPA051NSA_18 | Enteritidis |
| SAMN47165202 | 22EPA051NSA_19 | Enteritidis |
| SAMN47165203 | 22EPA051NSA_20 | Enteritidis |
| SAMN47165204 | 22EPA053NSA_1 | II salamae |
| SAMN47165205 | 22EPA053NSA_2 | II salamae |
| SAMN47165206 | 22EPA053NSA_3 | II salamae |
| SAMN47165207 | 22EPA053NSA_4 | II salamae |
| SAMN47165208 | 22EPA053NSA_5 | II salamae |
| SAMN47165209 | 22EPA053NSA_6 | II salamae |
| SAMN47165210 | 22EPA053NSA_7 | II salamae |
| SAMN47165211 | 22EPA053NSA_8 | II salamae |
| SAMN47165212 | 22EPA053NSA_9 | II salamae |
| SAMN47165213 | 22EPA053NSA_10 | II salamae |
| SAMN47165214 | 22EPA053NSA_11 | II salamae |
| SAMN47165215 | 22EPA053NSA_12 | II salamae |
| SAMN47165216 | 22EPA053NSA_13 | II salamae |
| SAMN47165217 | 22EPA053NSA_14 | II salamae |
| SAMN47165218 | 22EPA053NSA_15 | II salamae |
| SAMN47165219 | 22EPA053NSA_16 | II salamae |
| SAMN47165220 | 22EPA053NSA_17 | II salamae |
| SAMN47165221 | 22EPA053NSA_18 | II salamae |
| SAMN47165222 | 22EPA053NSA_19 | II salamae |
| SAMN47165223 | 22EPA053NSA_20 | II salamae |
| SAMN47165224 | 22EPA055NSA_1 | Anatum |
| SAMN47165225 | 22EPA055NSA_2 | Anatum |
| SAMN47165226 | 22EPA055NSA_3 | Anatum |
| SAMN47165227 | 22EPA055NSA_4 | Anatum |
| SAMN47165228 | 22EPA055NSA_5 | Anatum |
| SAMN47165229 | 22EPA055NSA_6 | Anatum |
| SAMN47165230 | 22EPA055NSA_7 | Anatum |
| SAMN47165231 | 22EPA055NSA_8 | Anatum |
| SAMN47165232 | 22EPA055NSA_9 | Anatum |
| SAMN47165233 | 22EPA055NSA_10 | Anatum |
| SAMN47165234 | 22EPA055NSA_11 | Anatum |
| SAMN47165235 | 22EPA055NSA_12 | Anatum |
| SAMN47165236 | 22EPA055NSA_13 | Anatum |

| | | |
|---|---|---|
| SAMN47165237 | 22EPA055NSA_14 | Anatum |
| SAMN47165238 | 22EPA055NSA_15 | Anatum |
| SAMN47165239 | 22EPA055NSA_16 | Anatum |
| SAMN47165240 | 22EPA055NSA_17 | Anatum |
| SAMN47165241 | 22EPA055NSA_18 | Anatum |
| SAMN47165242 | 22EPA055NSA_19 | Anatum |
| SAMN47165243 | 22EPA055NSA_20 | Anatum |
| SAMN47165244 | 22EPA058NSA_1 | Enteritidis |
| SAMN47165245 | 22EPA058NSA_2 | Enteritidis |
| SAMN47165246 | 22EPA058NSA_3 | Enteritidis |
| SAMN47165247 | 22EPA058NSA_4 | Enteritidis |
| SAMN47165248 | 22EPA058NSA_5 | Enteritidis |
| SAMN47165249 | 22EPA058NSA_6 | Enteritidis |
| SAMN47165250 | 22EPA058NSA_7 | Enteritidis |
| SAMN47165251 | 22EPA058NSA_8 | Enteritidis |
| SAMN47165252 | 22EPA058NSA_9 | Enteritidis |
| SAMN47165253 | 22EPA058NSA_10 | Enteritidis |
| SAMN47165254 | 22EPA058NSA_11 | Enteritidis |
| SAMN47165255 | 22EPA058NSA_12 | Enteritidis |
| SAMN47165256 | 22EPA058NSA_13 | Enteritidis |
| SAMN47165257 | 22EPA058NSA_14 | Enteritidis |
| SAMN47165258 | 22EPA058NSA_15 | Enteritidis |
| SAMN47165259 | 22EPA058NSA_16 | Enteritidis |
| SAMN47165260 | 22EPA058NSA_17 | Enteritidis |
| SAMN47165261 | 22EPA058NSA_18 | Enteritidis |
| SAMN47165262 | 22EPA058NSA_19 | Enteritidis |
| SAMN47165263 | 22EPA058NSA_20 | Enteritidis |
